# Transcriptome variation in human tissues revealed by long-read sequencing

**DOI:** 10.1101/2021.01.22.427687

**Authors:** Dafni A Glinos, Garrett Garborcauskas, Paul Hoffman, Nava Ehsan, Lihua Jiang, Alper Gokden, Xiaoguang Dai, Francois Aguet, Kathleen L. Brown, Kiran Garimella, Tera Bowers, Maura Costello, Kristin Ardlie, Ruiqi Jian, Nathan R Tucker, Patrick T Ellinor, Eoghan D Harrington, Hua Tang, Michael Snyder, Sissel Juul, Pejman Mohammadi, Daniel G MacArthur, Tuuli Lappalainen, Beryl Cummings

**Affiliations:** New York Genome Center, New York, NY, USA; Department of Systems Biology, Columbia University, New York, NY, USA; Medical and Population Genetics Program, The Broad Institute of MIT and Harvard; Department of Integrative Structural and Computational Biology, The Scripps Research Institute, La Jolla, CA, USA; Department of Genetics, Stanford University; Oxford Nanopore Technology, New York, NY, USA; Broad Institute of MIT and Harvard, Cambridge, MA, USA; Department of Biomedical Informatics, Columbia University, New York, New York, United States; Masonic Medical Research Institute, Utica, NY, USA; Cardiovascular Disease Initiative, The Broad Institute of Harvard and MIT, Cambridge, MA; Scripps Research Translational Institute, La Jolla, CA, USA; Analytical and Translation Genetics Unit, Massachusetts General Hospital, Boston, MA, USA; Centre for Population Genomics, Garvan Institute of Medical Research and Murdoch Children’s Research Institute

## Abstract

Regulation of transcript structure generates transcript diversity and plays an important role in human disease. The advent of long-read sequencing technologies offers the opportunity to study the role of genetic variation in transcript structure. In this paper, we present a large human long-read RNA-seq dataset using the Oxford Nanopore Technologies platform from 88 samples from GTEx tissues and cell lines, complementing the GTEx resource. We identified just under 100,000 new transcripts for annotated genes, and validated the protein expression of a similar proportion of novel and annotated transcripts. We developed a new computational package, LORALS, to analyze genetic effects of rare and common variants on the transcriptome via allele-specific analysis of long reads. We called allele-specific expression and transcript structure events, providing novel insights into the specific transcript alterations caused by common and rare genetic variants and highlighting the resolution gained from long-read data. We were able to perturb transcript structure upon knockdown of PTBP1, an RNA binding protein that mediates splicing, thereby finding genetic regulatory effects that are modified by the cellular environment. Finally, we use this dataset to enhance variant interpretation and study rare variants leading to aberrant splicing patterns.

## Main

Variation in transcript structure via RNA splicing and differences in the 5’ and 3’ untranslated regions (UTRs) is a key feature of gene regulation^1^. Disruption of transcript structure has a major role in human disease, with genetic variants associated with changes in splicing enriched in genome-wide associations for common diseases^2–4^ and implicated in many severe Mendelian diseases^5–7^. Common genetic variants affecting transcript structure can be mapped by transcript ratio and splicing quantitative trait locus (trQTL and sQTL) analyses that have further shown that genetic variants affecting gene expression levels and splicing tend to be distinct^8–10^. An orthogonal method to analyze genetic regulatory effects, allele-specific expression (ASE) analysis, has proven to be a highly sensitive method for studying rare genetic variants in *cis*^*11–13*^. However, the application of these approaches to short-read data relies on proxies for the full transcript structure and quantification, which are often inaccurate^14–18^. Furthermore, most metrics only attempt to quantify alternative splicing, leaving the role of UTRs obscure despite its demonstrated critical role in disease^19–21^. Long-read RNA sequencing technologies^22,23^ have now reached a mature stage, having already been used to study transcript structures^24,25^ and novel transcripts^26–28^, as well as early allele-specific analyses^29,30^. Allele-specific transcript structure (ASTS) analysis, enabled by long-read transcriptome data, could therefore provide important new information on how rare and common variants affect transcript structure and disease risk.

### Overview of dataset

Altogether, cDNA from 88 samples from 56 donors and 4 K562 cell line samples were sequenced on the MinION and GridION ONT platforms. Fibroblast cell lines were used to test the platform and to assess the direct-cDNA versus PCR-cDNA RNA-seq protocols (**Suppl. Figure 1A-C**). Since the primary purpose of this study was to study allelic events, which require high coverage, we prioritized depth and sequenced the remaining samples using the PCR-cDNA protocol. To evaluate the RNA isolation protocol, we used the K562 cell lines. The 88 GTEx samples included: 1) Assessment of replicability by three samples sequenced in duplicate and five samples in triplicate. Replicability was high (Spearman rho 0.87-0.95; **Suppl. Figure 1D**), leading us to merge the samples to increase depth; 2) The main dataset for analysis of transcriptome variation across tissues, consisting of 1-5 donors from 14 tissues; 3) Analysis of the effects of transcript perturbation by comparison of five GTEx fibroblast cell lines with and without PTBP1 RNA binding protein knockdown. Data were produced across two research centers (**Methods**; **Suppl. Table 1**). All the GTEx samples had Illumina TruSeq short-read RNA-seq data and 83 samples (51 donors) had whole genome sequencing data made available by the GTEx Consortium^4^.

Principal component analysis (PCA) and hierarchical clustering of samples based on transcript expression correlation showed tissue clustering (**Figure 1A,B** and **Suppl. Figure 1E**), similar to the GTEx consortium analysis of short-read RNA-seq data^4^. Gene and transcript quantifications from long-read data were highly concordant with those from Illumina RNA-seq (median R^2^=0.75 for genes and R^2^=0.57 for transcripts; **Figure 1C** and **Suppl. Figure 1F**). Genes and transcripts with low correlation were enriched for lower expression in ONT data, higher complexity genes and transcripts with multiple exons (**Suppl. Figure 2A-C**). We manually checked the read coverage of some of the genes that displayed low correlation, such as *PRELID1*, which is better captured by ONT, and *ARSB*, which displays 3’ bias (**Figure 1D**). Overall, cell lines with fresh RNA extracted with Trizol in Center 1 had lower 3’ bias than tissue samples and samples processed in Center 2 (**Figure 1E**).

**Figure 1:**
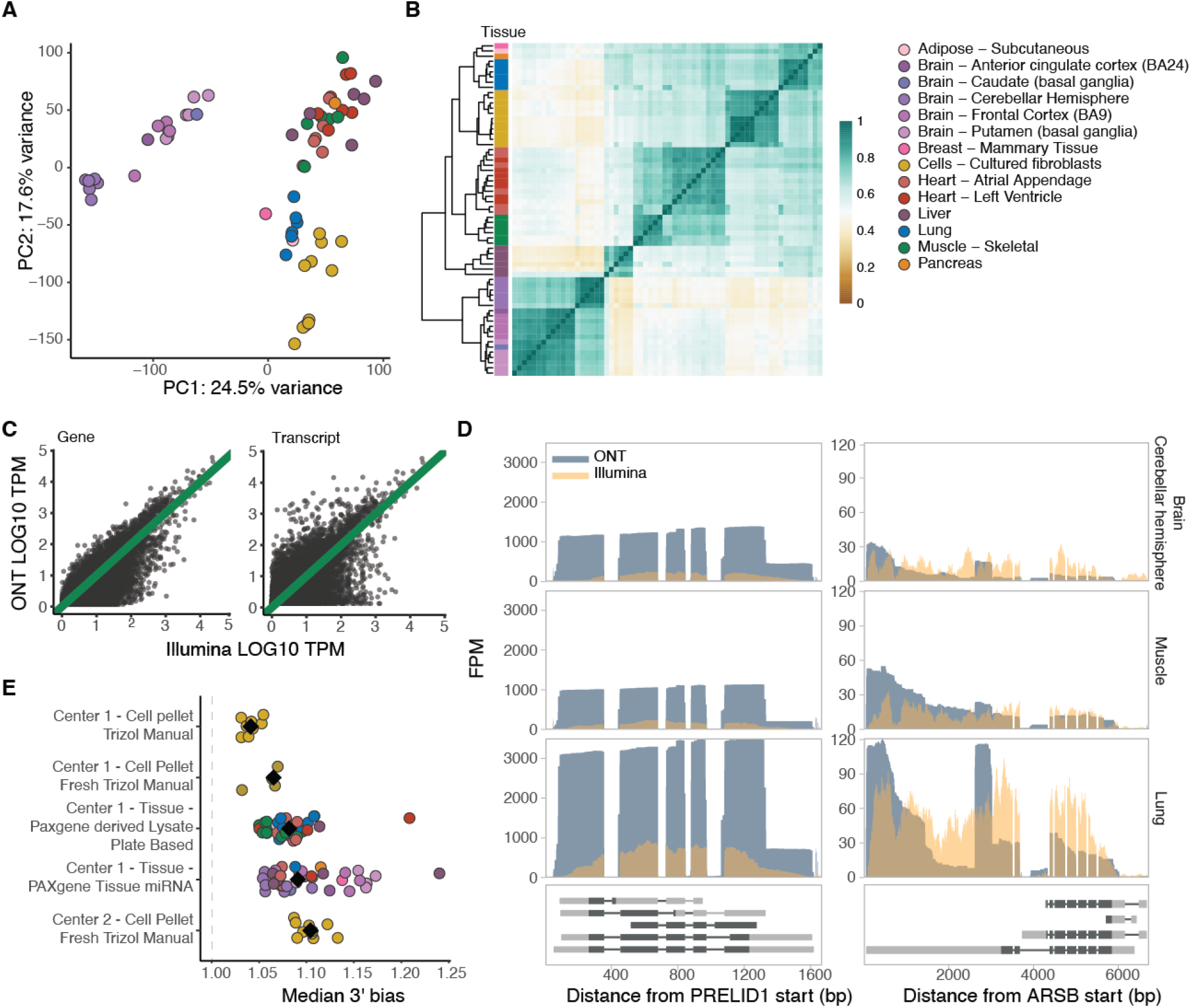
Overview and quality control of the dataset. **A)** Principal component analysis of samples with replicates merged, without K562 cell lines and without PTBP1 knockdown samples, based on GENCODE transcript expression (>3 TPM in >=5 samples). **B)** Hierarchical clustering of samples based on correlation of transcript expression (as in A), using Euclidean distance. **C)** Example of gene and transcript expression correlation between Illumina and ONT in the muscle tissue of GTEX-1LVA9. **D)** Two examples of genes displaying low correlation between ONT and Illumina. *PRELID1* was better captured by ONT than Illumina, while *ARSB* had 3’ bias when assayed by ONT. They are shown across three different tissues and all protein-coding transcripts are plotted below. FPM: Fragments per million. **E)** Median 3’ bias per sample, grouped by biospecimen type, RNA isolation method and sequencing center. Black diamonds indicate the median per group.

### Discovery of novel transcripts

We used FLAIR^31^ to quantify transcripts and identify novel ones, defined as transcripts with intron chains not matching with any transcript in GENCODE (v26) (**Methods**). We found 127,478 transcripts across 27,461 genes (**Suppl. Table 2**), of which 77% were novel (**Suppl. Figure 3A**). In most cases we quantified one, often already annotated, transcript for a gene, while more novel transcripts were discovered in genes with a high number of annotated transcripts (**Figure 2A**). Of the novel transcripts, 65,571 shared at least one splice junction with annotated transcripts (**Suppl. Figure 3B,C**) and 27,327 had intron retention, a significant enrichment compared to annotated ones (OR=3.8; **Figure 2B**). This suggests the presence of pre-mRNA despite carrying out a poly-A enrichment step. On the other hand, there was a modest depletion of exon skipping events in novel transcripts, suggesting they are well-represented in the existing annotations (OR=0.77; **Figure 2B**). We compared our findings with the 33,984 transcripts defined by Workman et al.^28^ based on GM12878 cell lines using ONT direct and cDNA RNA-sequencing, and detected 46% of the transcripts they identified, 4,584 of which were novel, providing further evidence to support the identified transcripts (**Suppl. Figure 3D**).

**Figure 2:**
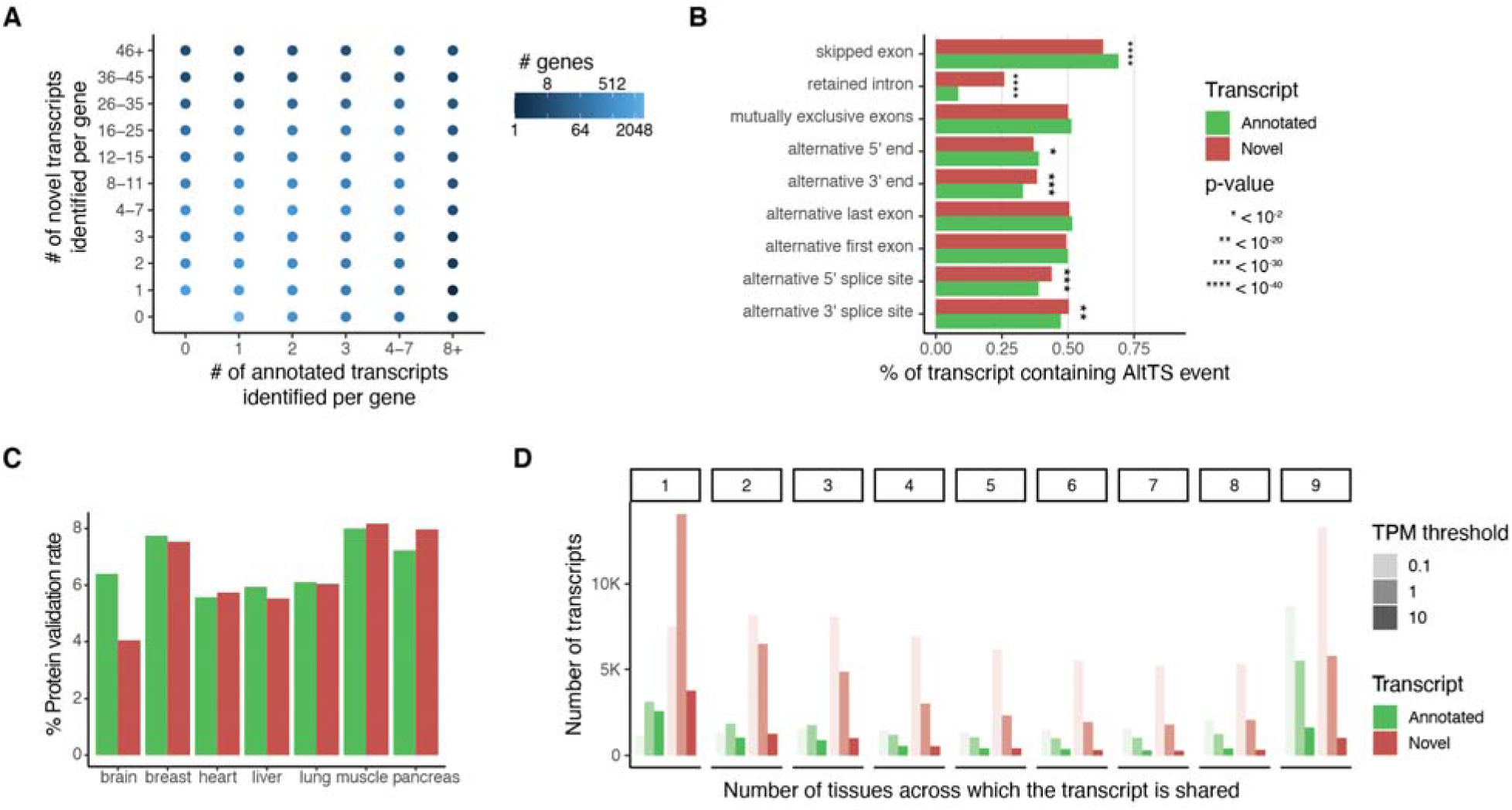
Discovery of new transcripts and comparison between tissues. **A)** Number of annotated and novel transcripts per gene quantified in our dataset. **B)** Proportion of alternative transcript structure (AltTS) events across all quantified transcripts, normalized per AltTS event. P-values were calculated using Fisher’s exact test. **C)** Percentage of validated transcripts at the protein level using mass spectrometry per primary tissue. **D)** Number of transcripts expressed across different TPM thresholds and classified based on how many tissues express the transcript at that level in at least two samples.

We validated our novel transcripts via proteome mass spectrometry data of 32 GTEx samples^32^. For most tissues we had assayed a similar number of samples using long-read RNA-seq and proteomics, with the exception of brain tissue, where additionally the sub-regions between the two assays did not match (**Suppl. Table 3**). We limited this analysis to 29,759 transcripts (61% of which were novel) expressed at ≥5 TPM in a sample per tissue and tested for matches in the predicted amino-acid chain. A comparable proportion of annotated and novel transcripts were validated in matched tissues (**Figure 2C**). In total, 2,397 unique transcripts were validated, of which 1,367 were novel, comparable to Jian et al^32^. Validated transcripts were enriched for exon skipping events, which was driven by novel transcripts, and both annotated and novel intron retention events showed lower validation rates (**Suppl. Figure 4A**,**B**). This depletion could be partially explained by nonsense-mediated decay or other post-transcriptional events depleting protein products rather than poor quality of the transcript annotations. For 201 genes we validated more than one transcript (445 total), with 294 transcripts being novel, often detecting tissue-specific protein transcript expression (**Suppl. Table 4** and **Suppl. Figure 4C**).

Novel transcripts resulted in clearer clustering of samples by tissue based on transcript expression correlations and PCA (**Suppl. Figure 5A**,**B**), suggesting that novel transcripts capture tissue-specific expression patterns. We therefore examined the gene and transcript expression across nine tissues with at least five samples. Highly-expressed novel transcripts were tissue-specific, with 55.5% expressed in a single tissue at >1 TPM (**Figure 2D**). This may explain their absence in existing annotations and highlights the potential for characterizing tissue-specific gene expression and regulation with long-read transcript analysis. We found thousands of transcripts exclusively expressed in a single tissue or having different transcript ratios across all nine tissues (**Suppl. Figure 6**). Of the novel transcripts exclusive to a tissue, the highest ratios were specific to the cerebellar hemisphere and the liver (41% and 19% respectively), concordantly with previous observation of high transcript diversity^33,34^.

### Allele-specific analysis

Allele-specific analysis captures *cis*-regulatory genetic effects on expression and transcript structure^8^. The expression of a gene or a transcript is quantified for each haplotype of a sample, separated based on the allele at a heterozygous site. Seventy-two of the long-read RNA-seq samples also had phased whole genome sequencing information from GTEx^4^, which allowed us to carry out allelic analysis. To address local alignment biases caused by sequencing errors adjacent to the variant sites of interest, we developed an alignment pipeline where two haplotype-specific references are created for each donor (**Suppl. Figure 7**). To perform allele-specific expression (ASE) and allele-specific transcript structure (ASTS) analysis, we developed a new software package, LORALS (Long-Read Allelic analysis). In addition to adopting mappability and genotyping error filters previously developed for short-read data^35^, we introduced flags addressing the higher error rate of long-read data (**Suppl. Figure 8**). Furthermore, we performed power calculations using simulated data to test how read counts, number of transcripts, and effect size affect ASTS detection power (**Suppl. Figure 9**).

Having established and optimized our pipeline, we performed the analysis using the FLAIR-aligned transcripts. Per sample, an average of 9% of genes analyzed for ASE or ASTS had a statistically significant event. To maximize power for generalizable insights, we analyzed all ASE (3,418 significant out of 34,255 across 6,370 unique genes) and ASTS events (321 significant out of 3,527 across 1,098 unique genes) combined across samples (**Suppl. Figure 10**). For 77% of genes analyzed for ASTS we quantified and tested the counts of 2 transcripts per gene, while the remaining ranged between 3 and 13.

Comparing the long-read ASE events to the ones reported for short-read GTEx v8 data^35^, we observed moderate concordance when looking at the p-values in short-read data using the long-read significant ASE events (π1 = 0.23) and vice-versa (π1 = 0.41) (**Suppl. Figure 11A**). Of the 354 events that were significant in both datasets, 84% had the same direction of effect (**Suppl. Figure 11B**). Differences were explained by low read depth and some variants being filtered out in one of the datasets (**Suppl. Figure 11C**), for example, 433 variants with significant ASE in long-read data were filtered in short-read data due to the mapping bias flag. Next, we sought to establish that ASE and ASTS recapitulate genetic regulatory effects of expression and splicing QTLs (eQTL and sQTL) mapped by GTEx^4^. Individuals who are heterozygous for a QTL lead SNP are expected to show increased allelic imbalance compared with those who are homozygous, and such significant enrichments were observed in the data (**Figure 3A**).

**Figure 3:**
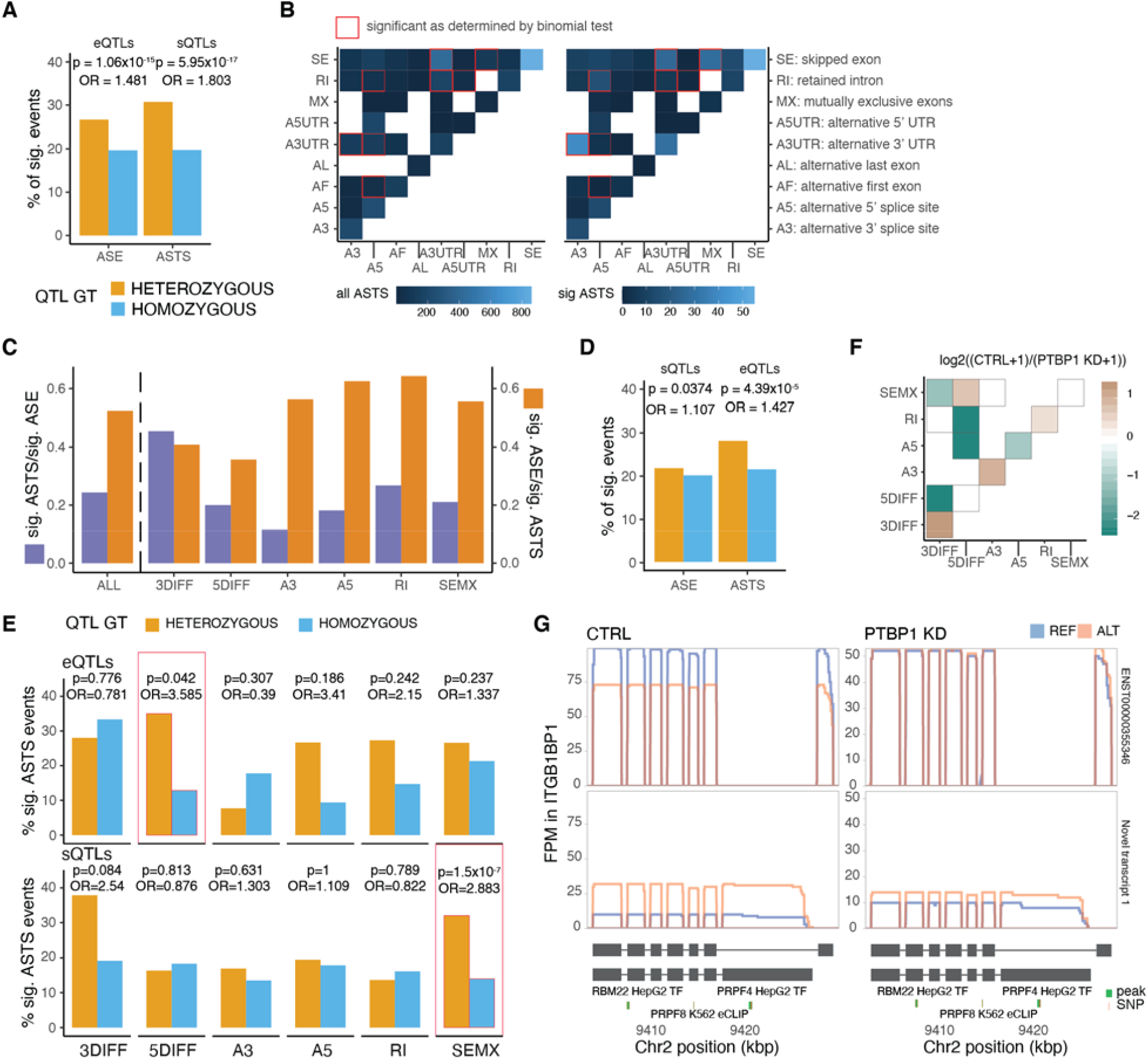
Allelic analysis of long-read data. **A)** Percentage of significant allele specific expression and transcript structure events for samples that are heterozygous or homozygous for a lead eQTL or sQTL variant for that gene, respectively. P-values from Fisher’s exact test. **B)** Co-occurrence of alternative transcript structure events within the transcripts used for ASTS analysis that are observed at least once per each event (or a single time for the diagonal) in a given gene. **C)** Sharing of ASE and ASTS events for all events, and stratified by AltTS event. **D)** Percentage of significant allele specific expression and transcript structure events for samples that are heterozygous or homozygous for a lead sQTL or eQTL variant for that gene, respectively. **E)** Percentage of significant ASTS for samples that are heterozygous or homozygous for a lead eQTL or sQTL variant for that gene, respectively, by type of event based on whether at least 50% of the differences in transcript can be assigned to that AltTS event. P-values from Fisher’s exact test. **F)** Changes in ASTS by PTPB1 knockdown, with the heatmap showing the co-occurrence alternative transcript structure events that are observed at least once per each event (or a single time for the diagonal) in a given gene. Color corresponds to the log2 ratio of the number of events found in the control over PTBP1 knockdown (KD) samples. **G)** *ITGBP1* gene transcript read pile-ups that display significant ASTS only in the control sample. Also shown are the transcript structures and the eCLIP and ChIP-seq for RNA binding proteins as assessed by ENCODE

Classification of alternative transcript structure (AltTS) changes enables better understanding of the nature of the ASTS events, and thus genetic variants affecting transcript structure. When considering each AltTS event alone, the most common was exon skipping, followed by alternative 3’ UTR events that were enriched for significant ASTS (**Suppl. Figure 12**). We then examined the combination of two types of AltTS events per gene (**Figure 3B**). We observed that certain event combinations occurred more commonly in significant compared to all ASTS events, for example the combination of alternative 3’ UTR with alternative 3’ splice sites (binomial test p-value = 1.6×10^−6^). On the other hand, there were combinations that were depleted from significant ASTS events, such as the combination of alternative 3’ UTR with retained introns (binomial test p-value = 7.6×10^−5^) or skipped exons (binomial test p-value = 1.2×10^−5^). This analysis highlights the prominent role of alternative UTR regions within the significant ASTS genes, missed in most sQTL mapping approaches.

In order to better understand the relationship between genetic effects on expression and transcript structure, we compared the ASE and ASTS events. We found that 203 of the 834 significant ASE genes displayed significant nominal p-values in ASTS (π1 = 0.13). This proportion was larger when looking at significant ASTS, where we found that 152 of the 320 genes displayed significant nominal p-values in ASE (π1 = 0.54; **Figure 3C**). This indicates that changes in transcript structure are often accompanied by changes in transcript levels, but less often the other way around. When repeating this analysis stratified by AltTS events we observed that an exception to this were ASTS events caused by alternative 3’ ends, where an equal proportion of events were ASE and ASTS.

Based on these observations, we examined sQTL-significant genes in ASE, where, as expected there was not a great difference between heterozygous and homozygous individuals (Fisher’s exact test p-value=0.0374). However, when looking at eQTLs, we observed that more heterozygous had significant ASTS compared to homozygous (Fisher’s exact test p-value=4.39×10^−5^; **Figure 3D**), indicating that genetically induced expression differences manifest in ASTS. In order to test the origin of this, we stratified the events by the AltTS events. We observed that the sQTLs were mostly manifesting in differences in exon skipping (32%; **Figure 3E**), as expected, while eQTLs were manifesting not only in total expression differences but also in transcript structure changes of the 5’ end of a gene (35%; **Figure 3E**). Differences in the 5’ end of a gene are therefore driving the capture of eQTLs in ASTS data, which would be normally missed by sQTL mapping.

This breakdown of events allows us to revisit existing sQTLs and find examples where ASTS data enables better understanding of the exact molecular events associated with the genetic variant, potentially contributing to diseases and traits (**Methods**). *DUSP13*, for example, is a gene specifically expressed in muscle, and has three sQTL intron excision phenotypes colocalizing with a single locus associated with body fat percentage. Multiple transcripts arise from this gene, but in both donors displaying ASTS we observed that the transcript ENST00000372700 lacking four middle exons was more highly-expressed from the risk allele (**Suppl. Figure 13A**). As further validation, GTEx short-read transcript ratios recapitulated this pattern (**Suppl. Figure 13B**). We were therefore able to pinpoint to the exact event leading to differences in transcript expression from the two alleles and potentially predisposing to high body fat percentage.

To test how ASTS captures changes in the effects of *cis*-regulatory variants due to perturbation of the cell’s splicing machinery, we knocked down PTBP1 RNA binding protein in five GTEx fibroblast cell lines. PTBP1 mediates exon skipping in pre-mRNAs and is involved in the 3′-end processing of mRNA. We therefore expected to see a disturbance of transcript expression as well as ASTS patterns for some genes upon siRNA knockdown. Indeed, we found 1,932 differentially expressed genes, 99.5% of which were validated with short-read data, and 1,742 differentially expressed transcripts. Exon skipping and alternative 3’ UTR events were enriched in transcripts upregulated in PTBP1 knockdown samples (**Suppl. Figure 14**).

We then compared allelic events in the knockdown and control samples (**Methods** and **Suppl. Figure 15A**,**B**), and observed an enrichment of condition-specific events in ASTS compared to ASE (Fisher’s exact test p-value = 0.0024; **Suppl. Figure 15C**), consistent with the fact that PTBP1 affects splicing and not gene expression at the allelic level. Control samples were enriched for ASTS with 3’ differences and 3’ alternative splice sites, while 5’ differences combined with 3’ differences, intron retention or 5’ alternative splice site ASTS events were enriched in the knockdown-specific ASTS (**Figure 3F**). This indicated that heterozygous genetic variants driving the ASTS in control samples lose their effect in the absence of PTBP1, and different transcript processing events take place. We hypothesized that ASTS disturbance upon PTBP1 knockdown is driven by heterozygous variants within RNA binding protein sites detectable in ChIP and eCLIP^36^ (**Suppl. Table 5**). We identified six control-specific ASTS events overlapping such sites. In *ITGBP1*, a donor has a heterozygous site within a PRPF4 binding site and ASTS that is strongly attenuated by PTPB1 knockdown (**Figure 3G**). A similar effect is seen in *PLAUR*, where there are two highly-expressed transcripts, one of which includes an exon skipping event (**Suppl. Figure 15D**). These analyses show how changes in the cellular environment altering splicing regulation can affect the molecular function of genetic variants.

### Rare variant interpretation

Finally, we evaluated the potential to better interpret rare variants with novel transcript annotations and ASTS data from long reads. We complemented the GENCODE v26 annotation with an additional 76,278 transcripts for protein-coding genes, and reannotated genetic variants from GTEx WGS data using VEP^37^ (**Methods**). The most severe consequence for a variant changed for 0.67% of all variants (**Suppl. Figure 16A**), 18,506 of which were coding (2.7% of coding variants). We used CADD scores as a proxy for the pathogenicity of a variant and as further support for validity of the re-classifications. We observed that variants reassigned to a more severe consequence had on average a higher CADD score than those that retained the same annotation (**Figure 4A**). An exception were variants previously annotated as 5’ UTR and reassigned as coding, but the already high CADD scores and selective constraint on 5’ UTR variants^38^ suggests that the UTR classification better reflects their functional impact. The higher CADD scores for variants reassigned as pathogenic provides independent evidence that our novel transcripts detect real biology and functional variants that may have been missed before. We therefore re-annotated ClinVar variants, resulting in the reassignment of 8,951 variants (1.2%). We observed that variants with benign clinical significance and no assertion criteria were reassigned at the highest rate (3.8%) while pathogenic variants reviewed by an expert panel were reassigned at the lowest rate (0.057%). Benign variants with low reviewer support were also reassigned at an order of magnitude higher rate than variants with pathogenic clinical significance (**Suppl. Figure 16B**). This provides an explanation for the conflicting reports of these variants and a potential pathogenic mechanism.

**Figure 5:**
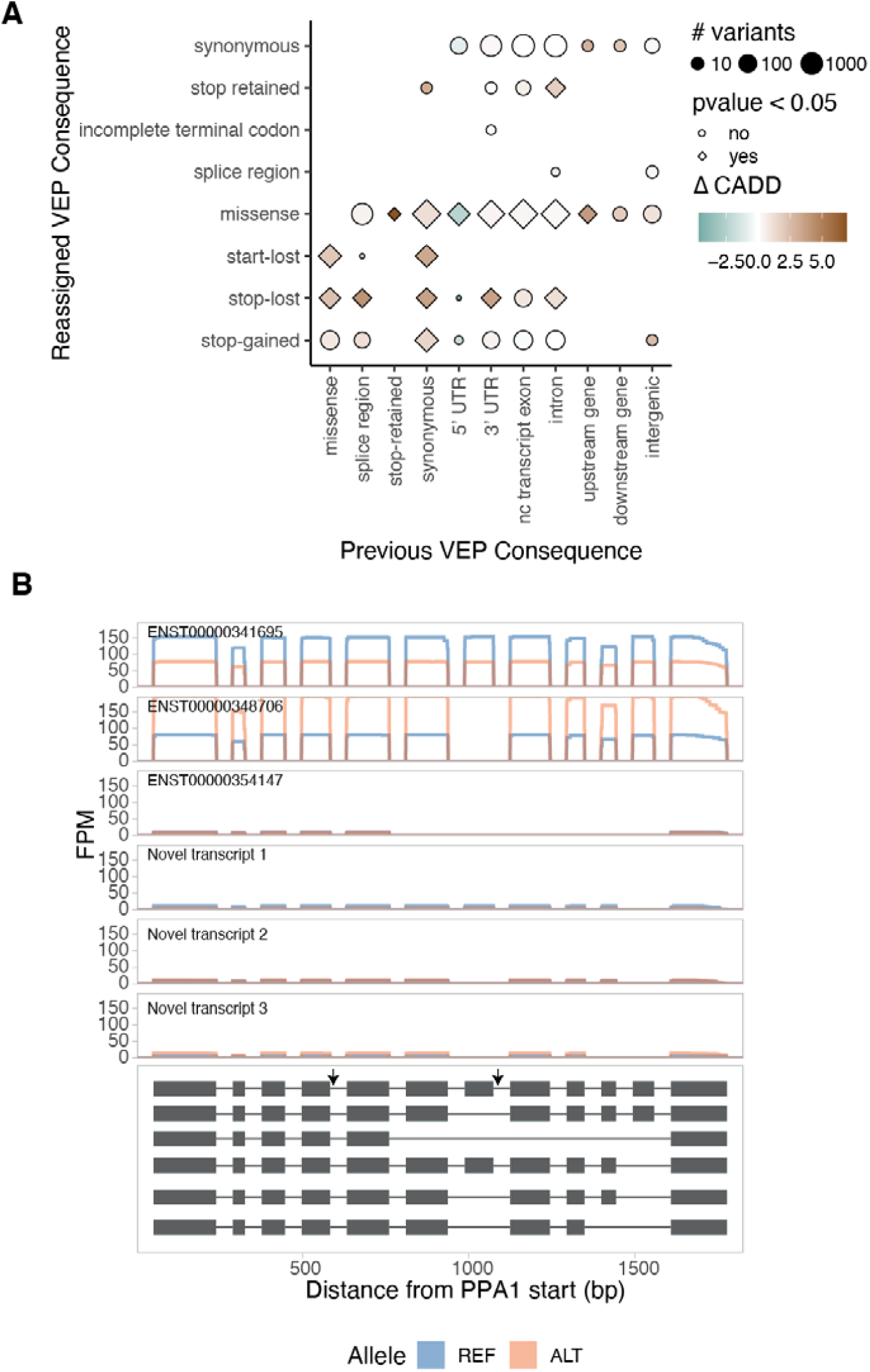
Variant interpretation through novel transcripts and allele-specific transcript structure analysis. **A)** Difference in the mean CADD score of variants that were reassigned to a more severe consequence when the GENCODE gene annotations were complemented with the novel FLAIR transcripts, compared to variants that retained their annotation (downsampled to a similar size). P-values from t-test. **B**) *PPA2* is an example of a gene with a rare heterozygous variant in a sample that is a GTEx splicing outlier and has significant ASTS, with read pileups, and grey arrows indicating the rare variants.

Long-read allelic data provides the opportunity to observe rare variants disrupting transcriptional regulation. GTEx has previously defined individuals that are extreme ASE, expression and splicing outliers, and shown that they are enriched for having rare genetic variants in the gene’s vicinity^13,39^. While our sample size is insufficient for analogous analysis of ASTS outliers, we tested the presence of rare (MAF<0.01) heterozygous variants within a 10kb window of each ASTS gene. Across all samples, missense variants were enriched for being in significant ASTS genes compared to all genes measured for ASTS (**Suppl. Figure 17A**,**B**). This indicates that ASTS can capture rare variant effects on transcript structure. Finally, we searched for specific examples where a rare variant is likely causing ASTS in our data (**Suppl. Table 6**). Out of ten genes where an individual has a rare heterozygous variant, is a splicing outlier as defined by GTEx, and has significant ASTS, we highlight two examples: *PPA2* has two intron variants chr4:105409456:G:A and chr4:105449015:G:A (MAF = 5.97×10^−4^ and 9.55×10^−3^) with the alternative allele having higher expression levels of transcript ENST00000348706 and lower expression of ENST00000341695 (**Figure 4B**) and *NDUFS4* (**Suppl. Figure 17C**,**D**).

## Discussion

In this study, we present the largest dataset of long-read RNA-seq to date, using material derived from cell lines and human tissues collected by the GTEx project. We identified 98,372 novel transcripts, which is higher than any other study^26–28^ likely due to our large sample size and tissue diversity, which is consistent with the high number of tissue-specific novel transcripts discovered. Supported by a high validation rate of the novel transcripts in high-throughput mass spectrometry proteome data^32^, our data makes an important contribution to human transcript annotations. Expanding long-read studies to further tissues and cell types, coupled with more extensive validation efforts, will enable better understanding of regulatory mechanisms of different types of transcript changes^26^, functionally distinct protein isoforms that different transcripts can give rise to^40^, and improved variant annotation, as demonstrated by our analysis.

Long reads provide the ability to map allelic effects over transcripts, instead of just expression^41^, thus providing the opportunity to analyze *cis* effects of genetic variants on transcripts. We developed LORALS, a toolkit for allelic analysis specific to long reads, taking into account various biases inherent to the technology. It is tunable and applicable to any long-read data, improving on previous work in this field^28,29^. We observed that the majority of ASTS events coincided with ASE, indicating that genetic effects on transcript usage rarely happen by reciprocally flipped transcript expression, but are typically accompanied by change in total expression levels which could happen for example via altered stability of specific transcripts^42^. However, the widespread co-occurrence of ASTS with ASE as well as eQTLs manifesting as ASTS are seemingly at odds with multiple QTL mapping studies that have established that expression and splicing are affected by distinct regulatory variants and processes^3,4,8^. The ability to distinguish the exact alternative transcript structure events in ASTS data allowed us to discover allele-specific 5’ differences as the cause of eQTLs manifesting in transcript structure changes, while expression and splicing are indeed highly independent. Given that promoter differences greatly affect gene expression levels and that most sQTL mapping methods do not capture variation in UTRs, this explains both the low overlap between causal variants of sQTLs and eQTLs and the overlap of ASTS with ASE and eQTLs.

These results reinforce the emerging understanding^20^ of the importance of analyzing the transcriptome not at the level of genes or imprecisely defined splicing, but rather with a detailed characterization of specific transcripts, their changes and combinations. These insights are readily captured by long-reads. Given the important role of genetic variants affecting transcript structure in disease risk^2–4,43,44^, we anticipate that a high-resolution characterization of the transcriptome with long-read data will be an important approach for the discovery of regulatory mechanisms of disease-associated variants.

## Data availability

Raw long read data generated as part of this manuscript are available in the GTEx v9 release under dbGAP accession number phs000424.v9. The GTEx WGS and Illumina short-read data are part of the GTEx v8 release phs000424.v8.

## Code availability

All original code used in the manuscript is released as part of a software package: https://github.com/LappalainenLab/lorals.

## Acknowledgements

We thank Michael Micorescu and Konstantinos Potamousis from the Oxford Nanopore Technologies commercial team for their help in generating the data.

## Authors contributions

DAG, TL and BC conceived and designed the project. DAG performed most of the data analysis. GG, AG and XD carried out the library preparation and sequencing. PH packaged the code. LJ, RJ, HT and MS provided and analyzed the data for the proteomic validation. PH and KB assisted in allelic expression analysis. KG carried out the base-calling. NE and PM performed power analyses and advised on analysis methods. AG carried out the PTBP1 knockdown. NRT and PE provided the CVD samples. TB and MC aided in the data generation. FA and KA coordinated the data release. FA, KA, EDH, SJ, DGM, BC and TL provided feedback on the study design and data analysis. NE, FA, NRT, EDH, SJ, PM and DGM provided feedback on the manuscript. EDH, SJ, DGM and TL supervised the work. DAG, TL and BC and wrote the manuscript with contributions from other authors. All authors read and approved the manuscript.

## References

1. Park, E., Pan, Z., Zhang, Z., Lin, L. & Xing, Y. The Expanding Landscape of Alternative Splicing Variation in Human Populations. Am. J. Hum. Genet. 102, 11–26 (2018).

2. Nicolae, D. L. et al. Trait-Associated SNPs Are More Likely to Be eQTLs: Annotation to Enhance Discovery from GWAS. PLoS Genet. 6, e1000888 (2010).

3. Li, Y. I. et al. RNA splicing is a primary link between genetic variation and disease. Science 352, 600–604 (2016).

4. GTEx Consortium. The GTEx Consortium atlas of genetic regulatory effects across human tissues. Science 369, 1318–1330 (2020).

5. Cummings, B. B. et al. Improving genetic diagnosis in Mendelian disease with transcriptome sequencing. Sci. Transl. Med. 9, (2017).

6. Kremer, L. S. et al. Genetic diagnosis of Mendelian disorders via RNA sequencing. Nat. Commun. 8, 15824 (2017).

7. Gonorazky, H. D. et al. Expanding the Boundaries of RNA Sequencing as a Diagnostic Tool for Rare Mendelian Disease. Am. J. Hum. Genet. 104, 466–483 (2019).

8. Lappalainen, T. et al. Transcriptome and genome sequencing uncovers functional variation in humans. Nature 501, 506 (2013).

9. Battle, A. et al. Characterizing the genetic basis of transcriptome diversity through RNA-sequencing of 922 individuals. Genome Res. 24, 14–24 (2014).

10. Li, Y. I. et al. Annotation-free quantification of RNA splicing using LeafCutter. Nat. Genet. 50, 151–158 (2018).

11. Rivas, M. A. et al. Human genomics. Effect of predicted protein-truncating genetic variants on the human transcriptome. Science 348, 666–669 (2015).

12. Smith, D. et al. A rare IL33 loss-of-function mutation reduces blood eosinophil counts and protects from asthma. PLoS Genet. 13, e1006659 (2017).

13. Mohammadi, P. et al. Genetic regulatory variation in populations informs transcriptome analysis in rare disease. Science 366, 351–356 (2019).

14. Trapnell, C. et al. Transcript assembly and quantification by RNA-Seq reveals unannotated transcripts and isoform switching during cell differentiation. Nat. Biotechnol. 28, 511–515 (2010).

15. Li, B. & Dewey, C. N. RSEM: accurate transcript quantification from RNA-Seq data with or without a reference genome. BMC Bioinformatics 12, 323 (2011).

16. Bray, N. L., Pimentel, H., Melsted, P. & Pachter, L. Erratum: Near-optimal probabilistic RNA-seq quantification. Nat. Biotechnol. 34, 888 (2016).

17. Teng, M. et al. A benchmark for RNA-seq quantification pipelines. Genome Biol. 17, 74 (2016).

18. Patro, R., Duggal, G., Love, M. I., Irizarry, R. A. & Kingsford, C. Salmon provides fast and bias-aware quantification of transcript expression. Nat. Methods 14, 417–419 (2017).

19. Pai, A. A. et al. Widespread shortening of 3’untranslated regions and increased exon inclusion are evolutionarily conserved features of innate immune responses to infection. PLoS Genet. 12, e1006338 (2016).

20. Alasoo, K. et al. Genetic effects on promoter usage are highly context-specific and contribute to complex traits. Elife 8, (2019).

21. Mittleman, B. E. et al. Alternative polyadenylation mediates genetic regulation of gene expression. Elife 9, (2020).

22. Sedlazeck, F. J., Lee, H., Darby, C. A. & Schatz, M. C. Piercing the dark matter: bioinformatics of long-range sequencing and mapping. Nat. Rev. Genet. 19, 329–346 (2018).

23. Amarasinghe, S. L. et al. Opportunities and challenges in long-read sequencing data analysis. Genome Biol. 21, 30 (2020).

24. Oikonomopoulos, S., Wang, Y. C., Djambazian, H., Badescu, D. & Ragoussis, J. Benchmarking of the Oxford Nanopore MinION sequencing for quantitative and qualitative assessment of cDNA populations. Sci. Rep. 6, 31602 (2016).

25. Weirather, J. L. et al. Comprehensive comparison of Pacific Biosciences and Oxford Nanopore Technologies and their applications to transcriptome analysis. F1000Res. 6, 100 (2017).

26. Anvar, S. Y. et al. Full-length mRNA sequencing uncovers a widespread coupling between transcription initiation and mRNA processing. Genome Biol. 19, 46 (2018).

27. Tardaguila, M. et al. SQANTI: extensive characterization of long-read transcript sequences for quality control in full-length transcriptome identification and quantification. Genome Res. (2018) doi:10.1101/gr.222976.117.

28. Workman, R. E. et al. Nanopore native RNA sequencing of a human poly(A) transcriptome. Nat. Methods 16, 1297–1305 (2019).

29. Tilgner, H., Grubert, F., Sharon, D. & Snyder, M. P. Defining a personal, allele-specific, and single-molecule long-read transcriptome. Proc. Natl. Acad. Sci. U. S. A. 111, 9869–9874 (2014).

30. Tilgner, H. et al. Comprehensive transcriptome analysis using synthetic long-read sequencing reveals molecular co-association of distant splicing events. Nat. Biotechnol. 33, 736–742 (2015).

31. Tang, A. D. et al. Full-length transcript characterization of SF3B1 mutation in chronic lymphocytic leukemia reveals downregulation of retained introns. Nat. Commun. 11, 1438 (2020).

32. Jiang, L. et al. A Quantitative Proteome Map of the Human Body. Cell 183, 269–283.e19 (2020).

33. Yeo, G., Holste, D., Kreiman, G. & Burge, C. B. Variation in alternative splicing across human tissues. Genome Biol. 5, R74 (2004).

34. Reyes, A. & Huber, W. Alternative start and termination sites of transcription drive most transcript isoform differences across human tissues. Nucleic Acids Res. 46, 582–592 (2018).

35. Castel, S. E. et al. A vast resource of allelic expression data spanning human tissues. Genome Biol. 21, 234 (2020).

36. Van Nostrand, E. L. et al. A large-scale binding and functional map of human RNA-binding proteins. Nature 583, 711–719 (2020).

37. McLaren, W. et al. The Ensembl Variant Effect Predictor. Genome Biol. 17, 122 (2016).

38. Whiffin, N. et al. Characterising the loss-of-function impact of 5’ untranslated region variants in 15,708 individuals. Nature Communications vol. 11 (2020).

39. Ferraro, N. M. et al. Transcriptomic signatures across human tissues identify functional rare genetic variation. Science vol. 369 eaaz5900 (2020).

40. Yang, X. et al. Widespread Expansion of Protein Interaction Capabilities by Alternative Splicing. Cell 164, 805–817 (2016).

41. Castel, S. E., Levy-Moonshine, A., Mohammadi, P., Banks, E. & Lappalainen, T. Tools and best practices for data processing in allelic expression analysis. Genome Biol. 16, 195 (2015).

42. Sibley, C. R. et al. Recursive splicing in long vertebrate genes. Nature 521, 371–375 (2015).

43. Scotti, M. M. & Swanson, M. S. RNA mis-splicing in disease. Nat. Rev. Genet. 17, 19–32 (2016).

44. Gandal, M. J. et al. Transcriptome-wide isoform-level dysregulation in ASD, schizophrenia, and bipolar disorder. Science 362, (2018).

