## Supplementary material for "Transcriptome variation in human tissues revealed by long-read sequencing": Supplemetary Material

### Methods

#### Fibroblasts cell culture and PTBP1 siRNA transfection

Patient derived immortalized fibroblast cell lines were cultured in DMEM media supplemented with 10% FBS and 1% Penicillin/Streptomycin (Corning). Transfections were performed 24h after initial seeding of 500,000 cells in 10cm dishes. Transfection mixtures were prepared with 6μg per dish of siRNA pools (Dharmacon SO-2720501G, SO-2703775G), Lipofectamine 2000 (Thermo Fisher), and Opti-MEM reduced serum media (Corning) according to proprietary guidelines. Mixtures were added to cell cultures containing reduced volumes of 5ml DMEM media for 6 hours before increasing volumes to 10ml with fresh media. Cells were harvested 96h after transfection.

#### SDS-PAGE and Western Blotting

Protein was extracted by boiling 75,000 cells at 95$^{o}$C for 5 minutes in 100μl 2 x Laemmli Sample Buffer (Bio-Rad) and 2-mercaptoethanol (5%) as a reducing agent. SDS-PAGE was run on 10% Mini-PROTEAN TGX gels (Bio-Rad) in Tris/Glycine/10%SDS Buffer. Proteins were transferred onto nitrocellulose membranes. 5% nonfat milk was used for blocking. Primary antibodies from mouse for PTBP1 (Thermo Fisher Scientific) and rabbit for GAPDH (Cell Signaling Technology) were incubated overnight at 4$^{o}$C. Secondary antibodies (LI-COR IRDye; donkey anti-mouse IgG polyclonal antibody (800CW; Size=100 μg) and donkey anti-rabbit IgG polyclonal antibody (680RD; Size=100 μg)) were incubated for 1h at room temperature (RT). Membranes were imaged on the Li-cor Odyssey CLx.

#### Generation of long-read RNA-seq data

Generally following the manufacturer's instructions, the below protocol was utilized with the following modifications.

**mRNA Isolation**

All tissue samples and six transfected fibroblast cell lines from the GTEx project were processed at the Broad Institute, along with the K562 cell lines. Five additional transfected fibroblast cell lines from GTEx, with and without PTBP1 knockout, were processed at the New York Genome Center (NYGC). For samples without biobanked RNA, total RNA was purified with the Direct-Zol RNA Miniprep kit (Zymo Research). Genomic DNA was digested using DNAse I (Ambion). The RNA was then repurified on columns to remove DNAse I and buffer. Quantifications and RIN scores were obtained on the BioAnalayzer (RNA kit, Agilent Technologies). Enrichment of mRNA was performed with Poly(dT) capture beads (Dynabeads, Thermo Fisher). For the samples processed at the Broad Institute, the volume of beads was scaled down to 25μL, and the isolation protocol was performed twice to ensure a pure mRNA product. Quantification of mRNA was obtained on the BioAnalyzer (RNA Pico kit, Agilent Technologies) and/or Qubit RNA HS assay (Thermofisher).

**Preparing input RNA (PCR) and reverse transcription**

Nine μL of the mRNA isolation was combined in a 0.2 mL PCR tube with 1μL VNP, and 1 μL 10 mM dNTPs (following the ONT PCS-109 protocol). The mixture was incubated at 65$^{o}$C for 5 minutes, and snap-cooled on a pre-chilled freezer block. In a separate tube, 4 μL 5x RT buffer, 1μL RNaseOUT (Invitrogen, cat #: 10777019), 1μL Nuclease-free water (ThermoFisher, cat #AM9937), and 2 μL Strand-switching primer (SSP) was combined. The strand-switching buffer was added to the snap-cooled, annealed mRNA and incubated at 42$^{o}$C for 2 minutes. Additionally the reads that contained over 8, 1 μL of Maxima H Minus Reverse Transcriptase (ThermoFisher, cat #EP0751) was added to each sample, which were then incubated for 90 minutes at 42$^{o}$C, followed by heat inactivation at 85$^{o}$C for 5 minutes. At NYGC, 1 uL of SuperScript IV reverse transcriptase (ThemoFisher, cat # 18091050) was used. The samples were incubated at 50 oC for 10 min and 42 oC for 10min, followed by 80 oC for 10min to inactivate.

**PCR**

Each RT sample was split into 4 PCR reactions in 0.2 mL PCR tubes, each containing 25 μL 2x LongAmp Taq Master Mix (NEB, M0287), 1.5 μL cDNA Primer, 18.5 μL Nuclease-free water, and 5 μL of the reverse-transcribed cDNA sample. The following cycling program in a standard PCR cycler was used: 30 seconds at 95oC, followed by 15 cycles of 95oC for 15 sec, followed by 62oC for 15 sec, followed by 65oC for 7 min. Final extension of 65oC for 6 minutes, followed by cooling down to 4oC. One µL Exonuclease I (NEB, M0293) was added directly to each PCR tube, and the reactions were incubated for 15 minutes at 37oC, and heat inactivated at 80oC for 15 minutes.

PerfeCTa SYBER Green FastMix (QuantaBio) was used for qPCR reactions, with 9µl of cDNA input and primer concentrations at 500nM. For PTBP1 the forward oligo sequence was 5'-ATT CTT TTC GGC GTC TAC GGT-3' and the reverse was 5'-TAT TGA ACA GGA TCT TCA CGC G-3' while for GAPDH the forward oligo sequence was 5'-CAT CCA TGA CAA CTT TGG TAT-3' and the reverse was 5'-CCA TCA CGC CAC AGT TTC-3'.

**Clean-up using AMPure XP beads**

To the 204µL of the PCR reactions in a clean 1.5 mL Eppendorf DNA LoBind we added 160 µL of resuspended AMPure XP beads (Beckman Coulter, A63880). The sample was mixed by pipetting gently 4x before being incubated at RT for 5 minutes. The sample was inverted gently 5x every 1-2 minutes. The samples were then spun down briefly and placed on a magnet to pellet the AMPure XP beads. The supernatant was carefully discarded and the pellet was washed 3x with 500 µL of fresh 70% EtOH. After drying, the pellet was resuspended in 204 µL of water and incubated at RT for 10 minutes, inverting every 2-3 minutes to keep the AMPure XP beads in suspension. The samples were placed on a magnet, and the supernatant was saved. Hundred twenty µL of AMPure XP beads were added to the 204 µL of supernatant and incubated for 5 minutes at RT, inverting gently 5x every 1-2 minutes.

The samples were spun down briefly and placed on a magnet to pellet the AMPure XP beads. The supernatant was removed and the pellet was washed 3x with 500 µL of fresh 70% EtOH. After the third wash, the samples were let to dry for 60 seconds. The pellet was resuspended in 12 µL of Elution Buffer (EB) and incubated at RT for 15-30 minutes. The microcentrifuge tube was flicked every 3-5 minutes to keep the AMPure XP beads in suspension.

After incubation, the samples were pelleted on a magnet and the supernatant was isolated and placed on ice. 1 µL of the supernatant was diluted into 9 µL of EB, and 2 µL of this 10x dilution was used to calculate the concentration, using HS DNA Qubit kit (ThermoFisher, Cat #: Q32851). 100 fmol of amplified cDNA was made up in 12 µL of EB.

**Adapter addition and loading flow cell**

One uL of the Rapid Adapter mix was added to the amplified cDNA library, and mixed by pipetting gently. The reaction was incubated for 5 minutes at RT and placed on ice until loading

The library was prepared for loading by mixing 37.5 µL Sequencing Buffer, 25.5 µL Loading Beads (mixed immediately before use by pipetting up and down), and 12 µL of the cDNA library.

After priming the flow cell and ensuring no bubbles, 75 µL of the prepared library was loaded onto the flow cell (flow cell type: FLO-MIN106) in a dropwise fashion. The library was run for 24-48 hours.

Optionally, 18-40 hours after the run began, 75 µL of a 1:1 mixture of SQB:water was added to the SpotON sample port to refuel the flow cell.

**Sequencing and basecalling**

Libraries were prepared with 300ng of input total RNA using the Illumina TruSeq kit and sequenced on the NextSeq 550 platform. Sequencing of mRNA samples was performed on the GridION X5 and MinION platform (Oxford Nanopore Technologies) for 48 hours.

To basecall the raw data we used ONT's Guppy tool (v2.3.7 for the samples processed at NYGC and v3.2.4 for the samples processed at the Broad Institute).

**Genome and transcriptome alignments**

We used minimap2 version 2.11[1] to align the reads to the GRCh38 human genome reference using -ax splice -uf -k14 parameters. We also aligned to the GENCODE v26 transcriptome using -ax map-ont parameters. We used NanoPlot [2] to calculate alignment statistics. We obtained a median of 6,304,430 raw reads per sample, of which on average 79% (s.d. 14%) aligned to the genome. The median read length was 709bp and 801bp for raw and aligned reads, respectively. We observed a higher median read length in samples sequenced using the direct-cDNA ONT protocol when compared to the PCR-cDNA protocol (t-test p-value = 0.022), at the expense of lower read depth (t-test p-value = $6.45 \times 10^{-3}$).

We used CollectRnaSeqMetrics from the Picard suite of tools to calculate 3' and 5' bias in our data (http://broadinstitute.github.io/picard/).

**Transcript detection and characterization**

We defined transcripts using FLAIR v1.4[3]. Four heart left ventricle samples from cardiovascular disease patients were included for the novel transcript calling. We used the samples that had been aligned to the genome and applied FLAIR-correct to correct misaligned splice sites using genome annotations. We merged all samples and ran FLAIR-collapse to generate a first-pass transcript set by grouping reads on their splice junctions chains and only keeping transcripts supported by at least 10 reads. Due to computational limitations, we split the first round alignments by chromosome at this step. We only kept reads with transcription start sites (TSS) that fell within promoter regions defined by taking a window 10bp upstream and 50bp downstream of the gene start site based on GENCODE v26 build and that spanned ≥80% of the transcript with ≥25 nucleotides coverage into the first and last exon. Reads that passed these filters were then realigned to the first-pass transcript set, retaining alignments with MAPQ >10.

We further filtered our transcript discovery set using TransDecoder software (https://github.com/TransDecoder/TransDecoder/) to remove transcripts with no ORFs. We integrated Pfam and Blast databases in this search, using the default parameters, in order to select the ORFs with the most functional coding potential. We removed transcripts where all open reading frames (ORF) were marked as being partial 3' and 5'. We further limited our discovery to transcripts encoding at least 100 amino acids long transcripts.

Transcripts were compared to GENCODE v26 and the ENCODE ONT release[4] using gffcompare[5]. Transcripts with exact intron chain-match were marked as annotated, while all others were marked as novel.

**Transcript quantification**

We used flair quantify[3] to quantify transcripts from all samples where reads had been aligned (1) GENCODE v26 and (2) the newly identified transcripts. Reads were normalised using transcripts per million normalisation and were filtered for transcripts expressed at least 5TPM in at least 3 samples prior to clustering analysis. Similarly, for the comparison between ONT and Illumina, reads were normalised using transcripts per million normalisation, filtered for protein-coding genes and limited to those with expression higher than 1TPM in both Illumina and ONT. Lowly correlated genes were defined by residual analysis of the Spearman correlations.

**Alternative transcript structure events definition**

We used SUPPA (v2.3) [6] to define alternative 3' splicing (A3), 5' splicing (A5), first exon (AF), last exon (AL), intron retention (RI), exon skipping (SE) and mutually exclusive exons (MX). We supplemented these annotations with alternative UTR regions, which for the purposes of this study were assumed to be the last exons. We used a window size of 10 nucleotides around splice sites, to allow for error.

**Protein validation of highly-expressed transcripts**

For the tissues assayed in the GTEx proteomics database[7] (heart, brain, liver, lung, muscle, pancreas and breast), we identified the transcripts expressed at higher than 5 TPM per sample. We used the output peptide fasta file from TransDecoder analysis in order to get the amino-acid sequence for each of the maintained transcripts. In total, 29,759 transcripts were maintained. To optimize our search-space, we grouped together brain samples from different regions as well as heart samples from different regions.

Raw files from the GTEx proteomics study[7] were first converted to mzXML files and then submitted to the Trans-Proteomic Pipeline (TPP) (http://tools.proteomecenter.org/wiki/index.php?title=Software:TPP) for database search. The Comet search engine was used for the database[8]. Mass tolerance of precursor ions was set to 10ppm and fragment ions was set to 1.0 amu. Up to two missed cleavages were allowed for trypsin digestion. Methionine oxidation was set to variable modification. Cysteine carbamidomethylation and peptide N-terminal and Lysine TMT modifications were set to be static modifications. After searches, peptides were filtered and scored by the PeptideProphet algorithm and proteins were scored afterwards using ProteinProphet [9]. Protein probability of 1 was used for the confident identification of the transcript.

**Differential transcript expression and transcript usage**

We used the nine tissues with at least five samples (brain cerebellar hemisphere, frontal cortex and putamen, cultured fibroblasts, atrial appendage and left ventricle from the heart, liver, lung and muscle). Differential expression was performed with DEseq2[10] pairwise using the Wald method and across all samples using the likelihood ratio test (LRT). We used replicates using the function collapseReplicates and within the design matrix we included the date of sequencing to correct for any biases. We used a cut-off for statistical significance at FDR=0.05. Differential transcript usage was performed with DRIMSeq[11]. Only the replicate with the highest read coverage was maintained in the analysis. All analysis was done in a pair-wise manner, with a cut-off for statistical significance at FDR=0.05.

Differential gene and transcript expression analysis between the control and PTBP1 knockdown samples were performed in the same way as above. For differential gene expression we used quantifications made based on  the GENCODE gene annotation, since each gene's differential gene expression status was validated using Illumina RNA-seq protocol on the same samples. For transcript differential expression we used the FLAIR transcripts.

**Allele-specific analysis**

**Alignment strategy**

We used the bcftools package to filter for only heterozygous variants per donor. We complemented the WGS and short-read RNA-seq phasing by long-read RNA-seq read phasing with HAPCUT2[12], run using all available RNA-seq libraries per subject. The haplotype phasing had been informed by the short-read RNA-seq data and we further switched the phase of a median of 2.2% of the heterozygous variants using the long-read data.

We generated a reference genome per haplotype of each donor and re-aligned the reads to each of the two references using the same parameters as described above. For each read, we retrieved the two MAPQ scores, and if different kept the one with the highest score; while the ties were randomly chosen between the two references. This approach led to a difference in alignment for on average 4.7% of the reads containing a heterozygous variant. It also allowed us to determine the source of the reference bias by looking at the start position of each read with each alignment strategy. Over 99% of the reads containing at least one heterozygous variant aligned at the exact same position, suggesting that the reference bias is due to local misalignment within the read, probably stemming from insertions/deletions adjacent to the variant of interest.

**Data acquisition**

SNP-level allele-specific data was generated using a software developed specifically for long-read data (LORALS). We flagged multi-mappability sites, sites that were part of the blacklist regions from ENCODE and monoallelic sites as determined by GTEx. Regions with multi-mapping reads were constructed using the alignability track from UCSC using a threshold of 0.1, meaning that a 100-kmer aligning to that site aligns to at least 5 other locations in the genome with up to 2 mismatches. Monoallelic sites were defined across all of their tissue for each sample, by testing whether there are no more reads supporting two alleles than would be expected from sequencing noise alone, indicating potential genotyping errors (FDR < 1%).

We introduced two ONT-specific flags, namely, the ratio of reference and alternative allele containing reads to the total read number for a site, which we set to greater than 80%, and the number of reads containing indels within a 10bp window of the heterozygous variant. This filter was determined by counting the number of matched base-pairs that were matched within the window and required at least 8 of them to not be INDELs. If at the site the proportion of indel containing was greater than 80% it was flagged. Additionally, the reads that contained over 8 INDELs within the window were filtered out. Finally, only variants that were covered by at least 20 reads were kept.

After filtering the flagged sites, we maintained a median of 73% (s.d. 4%) of the sites per sample, with the most stringent filter being the ratio of indel containing reads which removed 26% of the sites (s.d. 4%). For the variant sites that passed these filters we checked to which transcript each read was assigned to. We then created haplotype tables per gene across all of its transcripts. These tables were filtered for genes that had at least two transcripts, where each transcript has at least 10 reads.

We compared the reference ratios per gene and transcript across the samples for which we had either data from more than one tissue or which were sequenced in duplicates or triplicates. We observed a higher spearman correlation for samples from the same tissue (median R2 = 0.72 for ASE and R2 = 0.96 for ASTS) compared to samples from different tissues (median R2 = 0.65 for ASE and R2=0.83 for ASTS). We therefore merged the duplicate samples to increase our read depth.

**Statistical analysis, simulations and power analysis**

To analyze allele-specific analysis is based on the framework outlined in[13,14]. For a given gene and biallelic variant v, we define allelic expressions $e_0$, and $e_1$ as the sum of all transcripts produced from a gene located on the same chromosome copy as alleles each allele, respectively. For a given variant, we define log aFC as the expression originating from the alternative allele versus the reference allele (Eq. 1) and the reference ratio as the proportion of the reads originating from the reference allele over the total number of reads (Eq. 2)

$$s_{1,0} = log2 \frac{e_1}{e_0} \qquad \text{(Eq. 1)}$$

$$r_{1,0} = \frac{e_0}{e_0+e_1} \qquad \text{(Eq. 2)}$$

To test for statistical significant allele specific analysis, a binomial test was used to determine whether $r_{1,0}$is significantly different from the expected 0.5. Binomial test p-values were corrected for multiple hypothesis testing using the Benjamini-Hochberg procedure (FDR < 5%).

When testing for allele specific transcript structure, we performed power analysis to estimate the fraction of the cases where the distribution of transcript expression produced from the gene on the haplotypes were significantly different. Let $e_{hj}^{ti}$ be the allele-specific dosage for the transcript ($t_i$) from haplotype $h_j$. We denote $p_{hj}^{ti}$ as the allelic expression fraction of the transcript $t_i$, where $\sum_i^t = 1 \times p_{h_j}^{t_i} = 1$. The dependence of the two distributions $e_{h1}^{(t)}$ and $e_{h2}^{(t)}$ is determined by the chi-squared test ($x^2$).

The read counts, number of transcripts for each gene and the log aFC[14] are the factors that affect the power of the statistical test. Regarding the aFC factor, the maximum power happens at log aFC equal to zero, indicating equal expression in both haplotypes. Thus, for our analysis we assume that the log aFC is zero and statistical power is estimated to determine the dependency of allele-specific transcript structure analysis on the total coverage and transcript counts to detect an effect of a given size. The effect size is given by cohen's w, as defined in equation (Eq. 3)[15],

$$w = \sqrt{\frac{x^2}{N}} \qquad \text{(Eq. 3)}$$

This is applied on 2xm (m=number of transcripts) contingency table from $p_{h1}^{(t)}$ and $p_{h2}^{(t)}$ where, N is the total count table which in this case is 2. To give an idea of how the change in transcript ratios affects the magnitude of the effect size, the following $p_{h1}^{(t)}$ and $p_{h2}^{(t)}$ pairs are presented in all of which w = 0.3 (interpreted as medium effect size).

| | | | | | | | | | | | |
|---|---|---|---|---|---|---|---|---|---|---|---|
| t=2 | $p_{h_1}^{(t)}$ | .5 | .5 | | | | | | | | |
| | $p_{h_2}^{(t)}$ | .8 | .2 | | | | | | | | |
| t=3 | $p_{h_1}^{(t)}$ | .33 | .33 | .34 | | | | | | | |
| | $p_{h_2}^{(t)}$ | .33 | .57 | .1 | | | | | | | |
| t=4 | $p_{h_1}^{(t)}$ | .25 | .25 | .25 | .25 | | | | | | |
| | $p_{h_2}^{(t)}$ | .25 | .25 | .45 | .05 | | | | | | |
| t=5 | $p_{h_1}^{(t)}$ | .2 | .2 | .2 | .2 | .2 | | | | | |
| | $p_{h_2}^{(t)}$ | .2 | .2 | .2 | .38 | .02 | | | | | |
| t=10 | $p_{h_1}^{(t)}$ | .1 | .1 | .1 | .1 | .1 | .1 | .1 | .1 | .1 | .1 |
| | $p_{h_2}^{(t)}$ | .1 | .1 | .1 | .1 | .1 | .1 | .1 | .26 | .03 | .01 |

To perform the power estimation, the simulated allelic expressions $e_{h1}^{(t)}$ and $e_{h2}^{(t)}$ are produced from a multinomial distribution of two normalized random vectors $p_{h1}^{(t)}$ and $p_{h2}^{(t)}$ that specify the effect size of interest. The significant difference of $e_{h1}^{(t)}$ and $e_{h2}^{(t)}$ is determined by the chi-squared test (nominal p-value < 0.01). Power estimation based on simulated data for a set of read counts and number of transcripts for the effect sizes of 0.3 and 0.5, is calculated. The effect size is rounded for one digit. In order to detect ASTS with effect size 0.5 with 60% power, assuming aFC=0, total read coverage of 36 was required. For effect size 0.3, at least 100 reads were needed.

To analyze the allele-specific transcript structure using GTEx long-read sequencing data, p-values are derived from chi-squared test for allelic transcript expressions for all genes and all individuals. To filter genes with low expressed transcripts, we considered those genes that have more than one transcript which at least one of the two haplotypes carrying REF or ALT allele have at least 10 reads and in case multiple variants associated to a gene the one with the highest total coverage is selected for the analysis. P-values are corrected for multiple hypothesis testing using the Benjamini-Hochberg procedure (FDR < 5%).

**Comparison across datasets**

For the comparison between two datasets significant results we used pi1 statistic setting lambda between 0 and 0.8 in increments of 0.001. (http://github.com/jdstorey/qvalue). For all pi1 calculations we only used genes that could be captured in both datasets. Comparison to GTEx ASE events obtained by Illumina were done using the SNV-level read counts, annotated using the GENCODE annotations, for continuity. The datasets were merged and the variant with the highest read count across both methods was selected per gene across samples.

**Colocalization analysis**

We mined all colocalization results between GTEx sQTLs and 5,586 GWAS traits[16] and filtered for loci with rcp >0.5 and removed the HLA region. We then mapped each sQTL to its corresponding gene and overlapped that gene set with the significant ASTS genes per tissue. For the overlapping genes we verified that the lead sQTL used for colocalization was a heterozygous variant in the donor for which we had ASTS data. This strict filtering resulted in three genes *SRP14*, *DUSP13* and *ELP5*. After further investigation we removed *SRP14* because its signal seemed to originate from expression rather than structure.

**Combinatorial allele-specific analysis in control and PTBP1 KD samples**

For each donor the control and the knockdown samples were processed together, and the most highly covered variants using both samples were selected per gene. Specific allelic events per condition were defined using an FDR threshold of 0.05.

We downloaded all eCLIP (bed narrowPeak) and ChIP-seq (bed narrowPeak, IDR thresholded peaks) RNA protein binding data in GRCh38[17]. All peaks were overlapped with the heterozygous variants per donor using bedtools intersect[18]. Finally, the maintained peaks were annotated to the nearest gene using a 10kb window around each gene.

#### Annotation of variant consequences

Annotation of protein-coding regions was generated by running Ensembl VEP (version 100.2) with the --most_severe flag on the GTEx v8 release. We did two rounds of annotation, the first one using protein-coding genes from the GENCODE v26 GTF and the second one by supplementing this annotation with newly identified FLAIR transcripts for these genes. We predicted the productivity of each transcript using flair predictProductivity.py (v1.4)[3] using only the longest ORF for each transcript. The frame of each transcript was corrected using genomeTools (v1.6.1)[19]. Transcripts were classified as protein-coding if both a start and a stop codon were found, nonsense-mediated-decay if a premature termination codon was found, or as a processed transcript if there was no stop codon. Finally, the gene coordinates were extended if one of the transcripts was found to be outside them. This led to 76,278 transcripts added.

CADD scores for all annotated variants were obtained using the v1.5 release[20]. We compared the CADD scores between the reassigned and the non-reassigned variants (down-sampled to match the size of the total number of reassigned variants per consequence group). We then used a t-test to compare the means of the two groups.

#### Rare variant analysis

We extracted all heterozygous variants within a 10kb window around each gene assessed for ASTS in a donor specific manner. Variants were filtered for MAF <0.01 and the worst consequence was maintained per variant. We found a median of four rare variants per gene. We observed that 50% of genes across all samples had at least one rare variant. We calculated enrichment using a binomial test, setting all variants as background.

### Supplementary Tables

**Supplementary Table 1**: Samples sequencing metadata.

**Supplementary Table 2**: Counts of flair transcripts per gffcompare category.

**Supplementary Table 3**: Number of samples per tissue used for protein validation, in long-read RNA-seq and in mass spectrometry data.

**Supplementary Figure 4**: Highly expressed transcripts validated by mass spectrometry proteome data, where more than one transcript was observed for a single gene.

**Supplementary Table 5**: Number of sQTLs colocalizing with GWAS overlap with ASTS genes.

**Supplementary Table 6**: Heterozygous variants overlapping eCLIP and ChIP-seq peaks from the ENCODE database of RNA-binding proteins within a 10kb window of ASTS significant genes.

**Supplementary Table 7**: Rare heterozygous variants within 10kb of ASTS significant genes that were also observed to be splicing outliers.

### Supplementary Figures

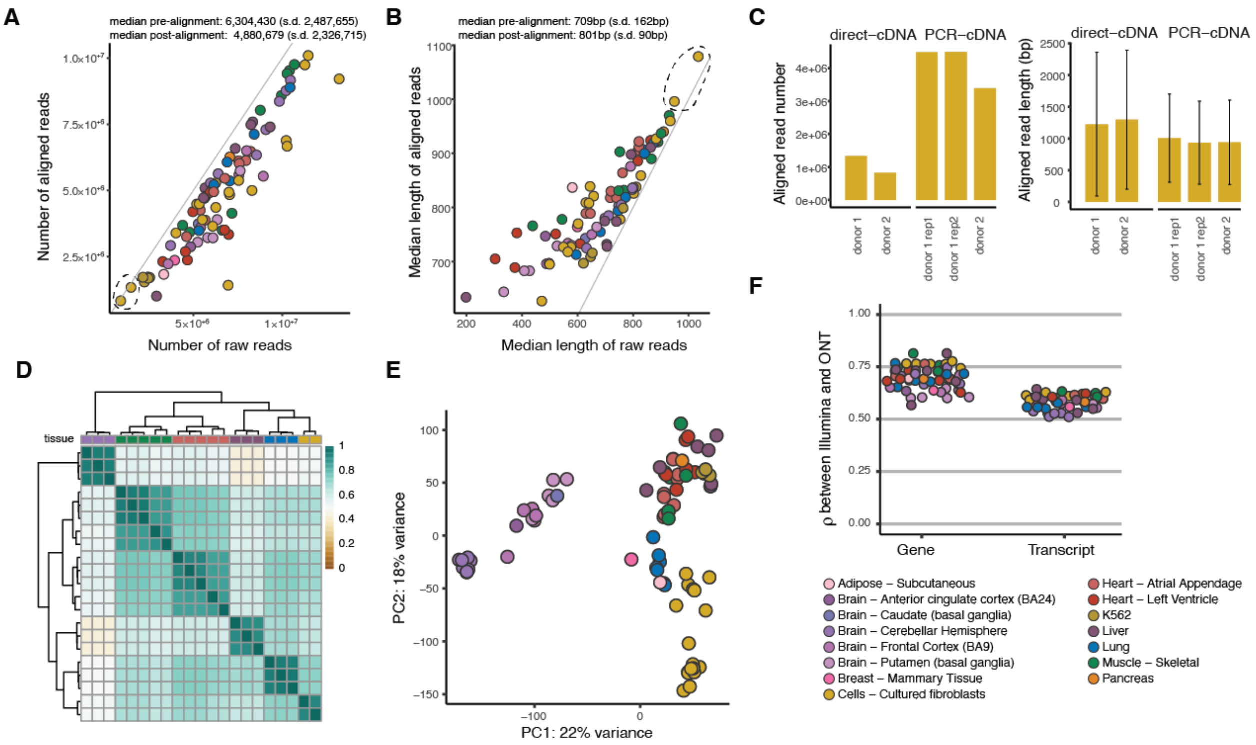


**Supplementary Figure 1: A)** Number and **B)** median length of raw and aligned reads per sample. **C)** Read number and read length in two fibroblast cell line samples (one of which was sequenced in replicate) that were sequenced using both the direct-cDNA and the PCR-cDNA protocol. Error bars: standard deviation from the mean. **D)** Hierarchical clustering using Euclidean distance for replicate samples aligned to GENCODE for transcripts with expression above 3 TPM in at least 5 samples. **E)** Principal component analysis using 88 samples aligned to GENCODE (v26) for transcripts with expression above 3 TPM in at least 5 samples. **F)** Correlation between the transcriptome of each sample quantified by ONT and by Illumina sequencing technologies.

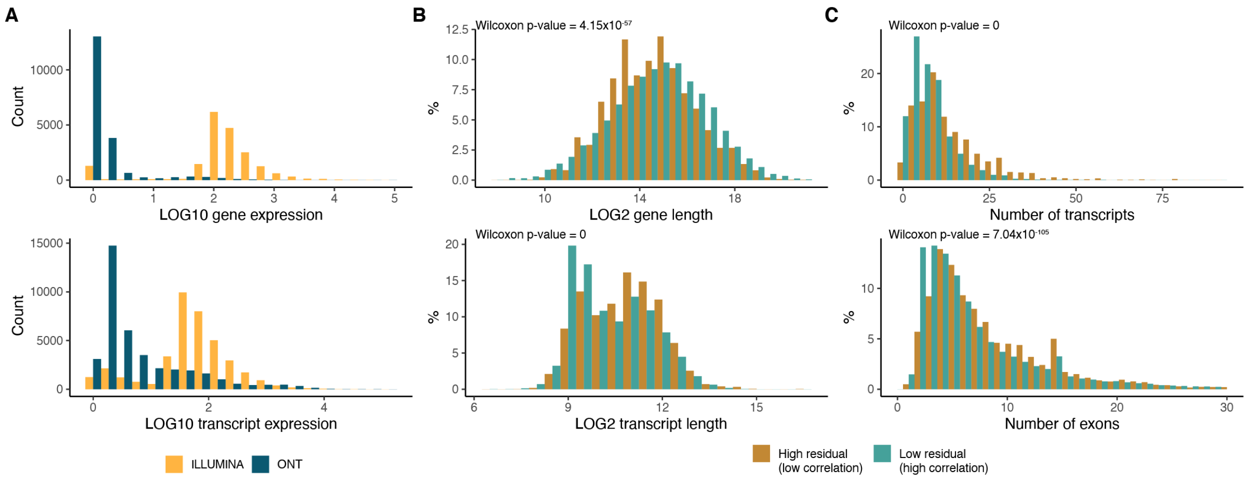


**Supplementary Figure 2**: **A)** Normalized gene and transcript expression for high residual (|residual| > 1) genes and transcripts retrieved from the Spearman correlation analysis. **B)** Characteristics of genes and **C)** transcripts with high or low residuals with respect to gene/transcript length, number of transcripts per gene and number of exons per transcript. The two groups' means were compared using Wilcoxon's test.

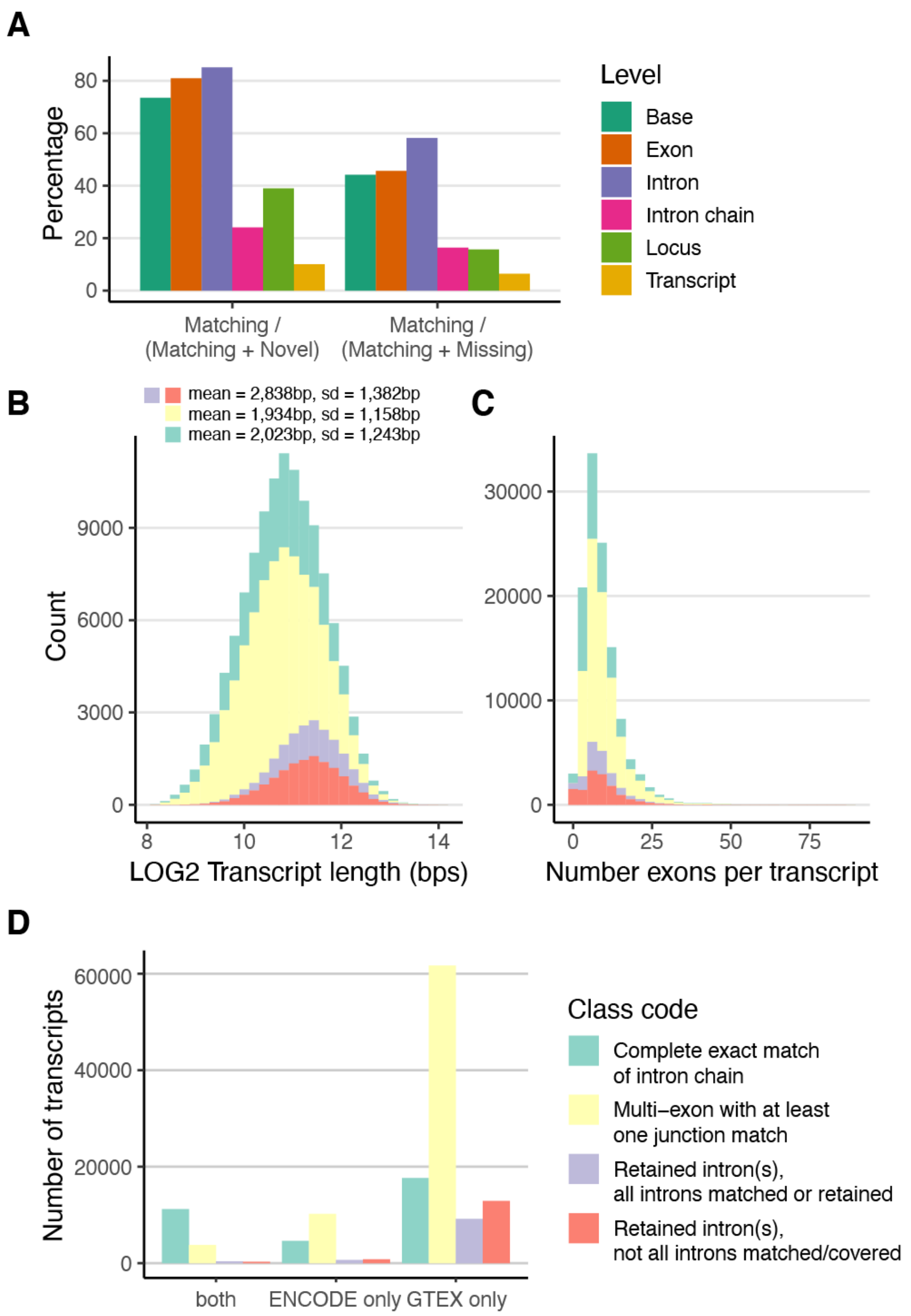


**Supplementary Figure 3**: **A)** FLAIR transcripts comparison to GENCODE with respect to different genomic levels. The difference between intron chain and transcript is that the former only looks at matching the intron boundaries, therefore allowing variation in the UTR regions. **B)** Transcript length and **C)** number of exons per transcript classified by comparison to GENCODE. **D)** Number of overlapping transcripts between the ones identified in this paper and the ones released by ENCODE.

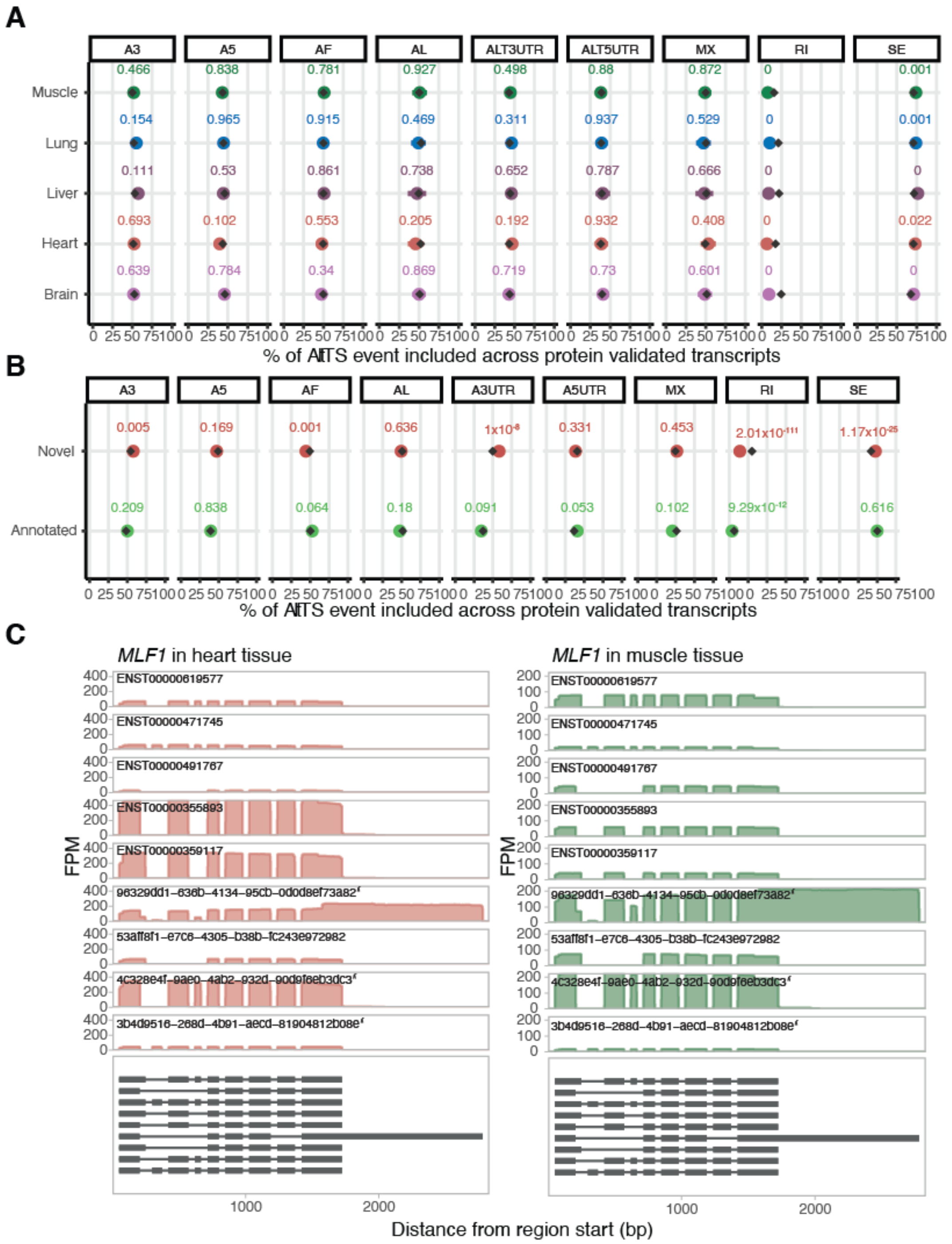


**Supplementary Figure 4**: Proportion of highly-expressed transcripts that were validated by mass spectrometry **A)** per tissue or **B)** per novel or annotated, split according to the AltST event that they belong (a transcript can be assigned to more than one category). Enrichment was calculated using a binomial test against all highly-expressed transcripts sent for validation. **C)** MLF1 is an example of a gene with multiple highly-expressed transcripts across both muscle and heart tissues with different transcripts validated in each (indicated by an asterisk*). A3: alternative 3' splice site; A5: alternative 5' splice site; AF: alternative first exon; AL: alternative last exon; A3UTR: alternative 3' end; A5UTR: alternative 5' end; MX: mutually exclusive exons; RI: retained intron; SE: skipped exon.

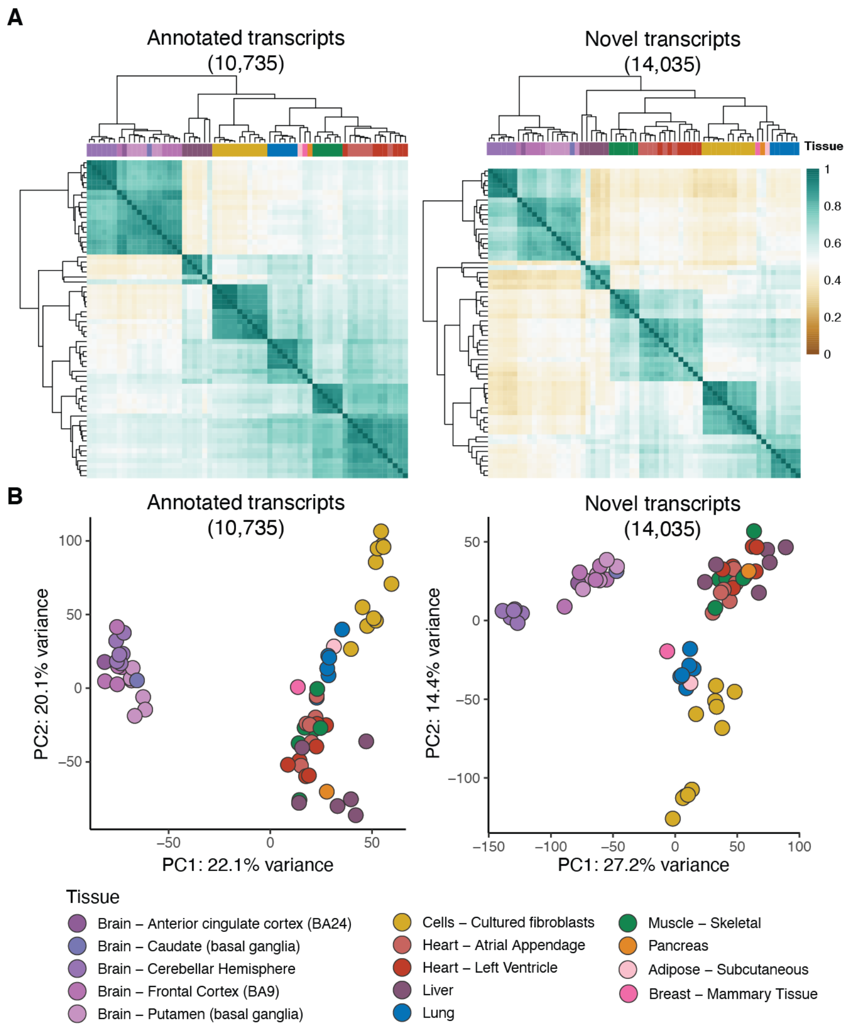


355

**Supplementary Figure 5**: **A)** Hierarchical clustering using euclidean distance and **B)** principal component analysis for selected samples aligned to GENCODE for transcripts with expression above 5 TPM in at least 3 samples separated based on whether they are novel or not.

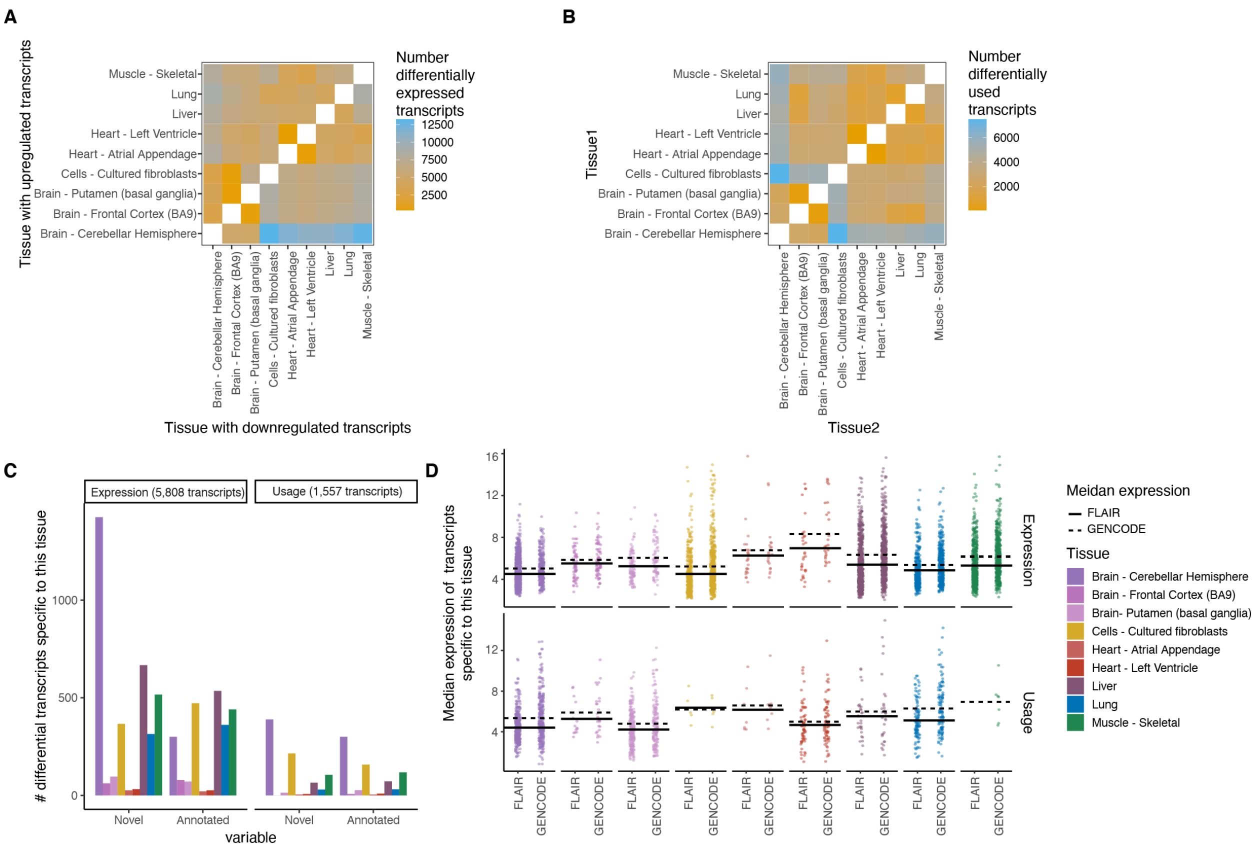


**Supplementary Figure 6**: Heatmap of number of differentially **A)** upregulated transcripts and **B)** used transcripts (FDR ≤ 0.05) in pairwise comparison of tissues with at least five samples. In differential expression analysis we identify up- or down-regulated transcripts per pairwise comparison (asymmetrical heatmap) while in the differential transcript usage analysis there is no direction of effect (symmetrical heatmap). **C)** Number of differentially expressed or used transcripts that were specific to that tissue. **D)** Median gene expression across all transcripts that are specific to a tissue.

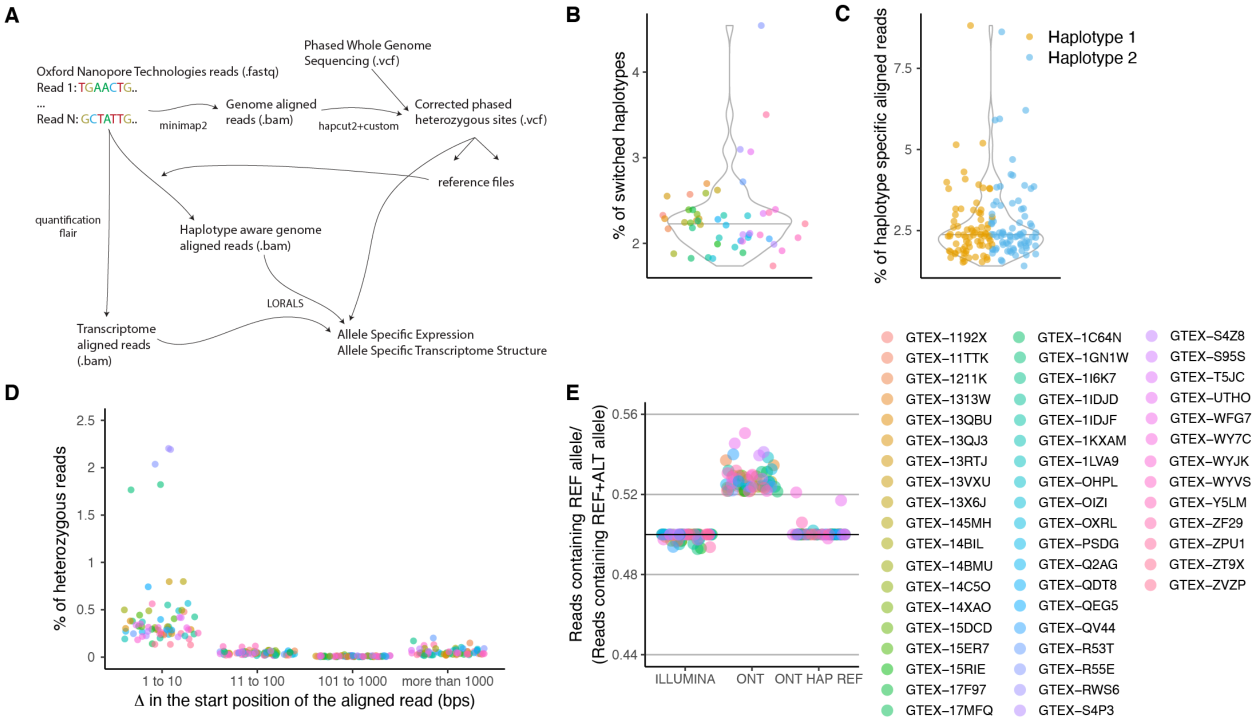


**Supplementary Figure 7**: **A)** Pipeline for allele-specific analysis. Raw long-reads are first aligned to the genome using minimap2. This alignment is used to correct the phase of some of the heterozygous variants on the whole genome sequencing vcf. This new file is then used to generate personalized genome reference files against which the raw reads are again aligned using minimap2. The raw reads are also aligned to the transcriptome using minimap2. The VCF file along with the genome aligned reads and the transcriptome aligned reads are then fed into LORALS for allelic analysis. **B)** Percentage of switched haplotypes per donor informed by the long-read data. For this all samples from the same donor were merged to harmonize the files. **C)** Percentage of haplotype specific reads calculated as reads having a higher mapping score when using a personalized genome reference. **D)** Delta calculated as the difference in the start position of the aligned read between the genome aligned and the personalized genome aligned reads. Not shown are the reads that had Delta = 0. **E)** Reference ratio for the samples present in this study sequenced using Illumina technology and ONT technology aligned with two different approaches.

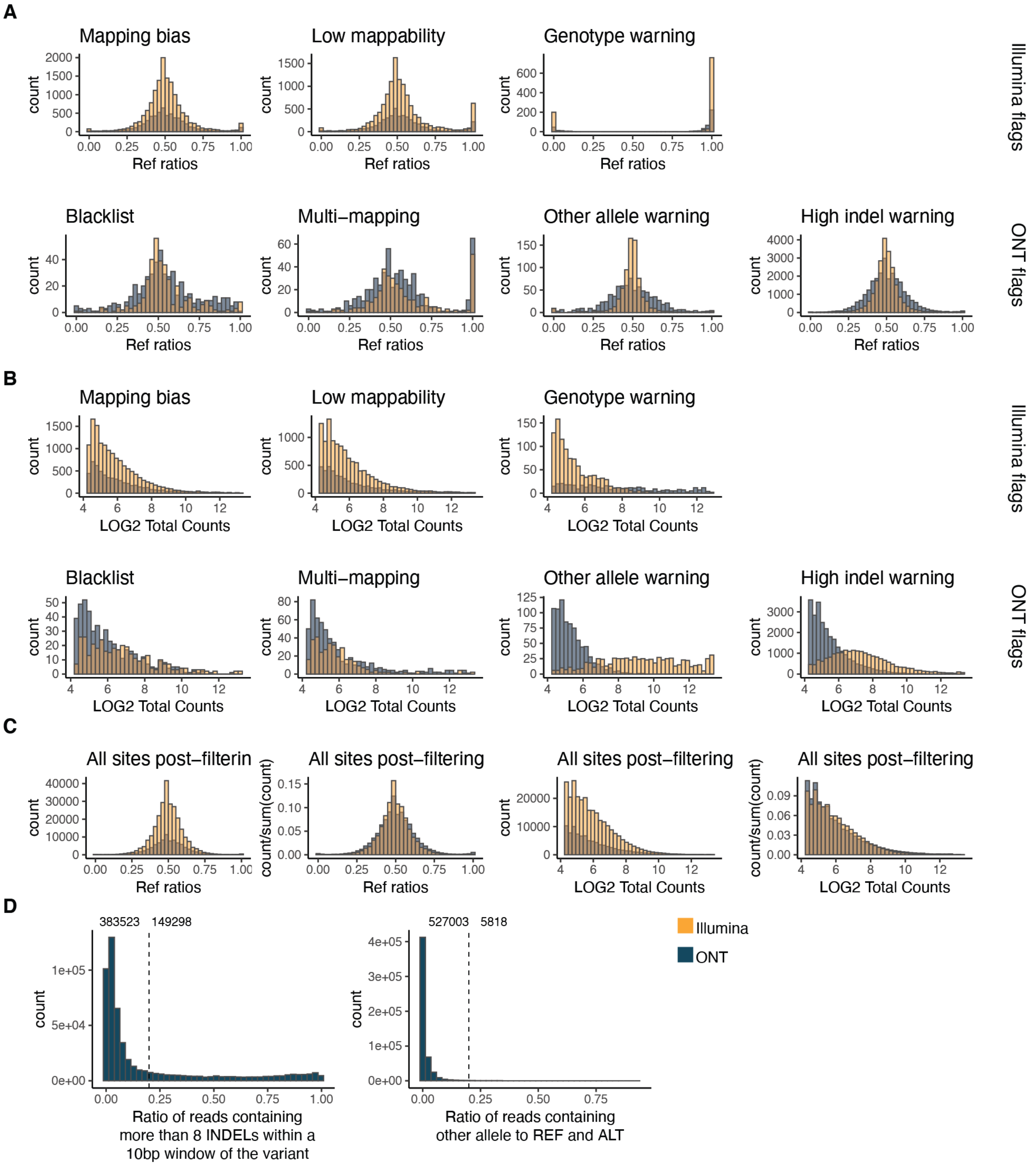


**Supplementary Figure 8: A)** Reference ratio and **B)** normalised reads counts across different Illumina and ONT flags for both of these sequencing technologies. Mapping bias: mapping bias in simulations; Low mappability: low-mappability regions (75-mer mappability < 1 based on 75mer alignments with up to two mismatches based on the pipeline for ENCODE tracks and available on the GTEx portal); Genotype warning: no more reads supporting two alleles than would be expected from sequencing noise alone, indicating potential genotyping errors (FDR < 1%); Blacklist: ENCODE blacklist. Multi-mapping: regions with multi-mapping reads constructed using the alignability track from UCSC using a threshold of 0.1 (so that a 100kmer aligning to that site aligns to at least 5 other locations in the genome with up to 2 mismatches); Other allele warning: regions where the proportion of ref or alt containing reads is lower than 0.8; High indel warning: sited where the proportion of non-indel containing is lower than 0.8. **C)** Reference ratios and normalised reads counts of all kept sites across Illumina and ONT sequencing technologies. **D)** Distribution of the high indel warning ratios and the other allele ratios across all samples.

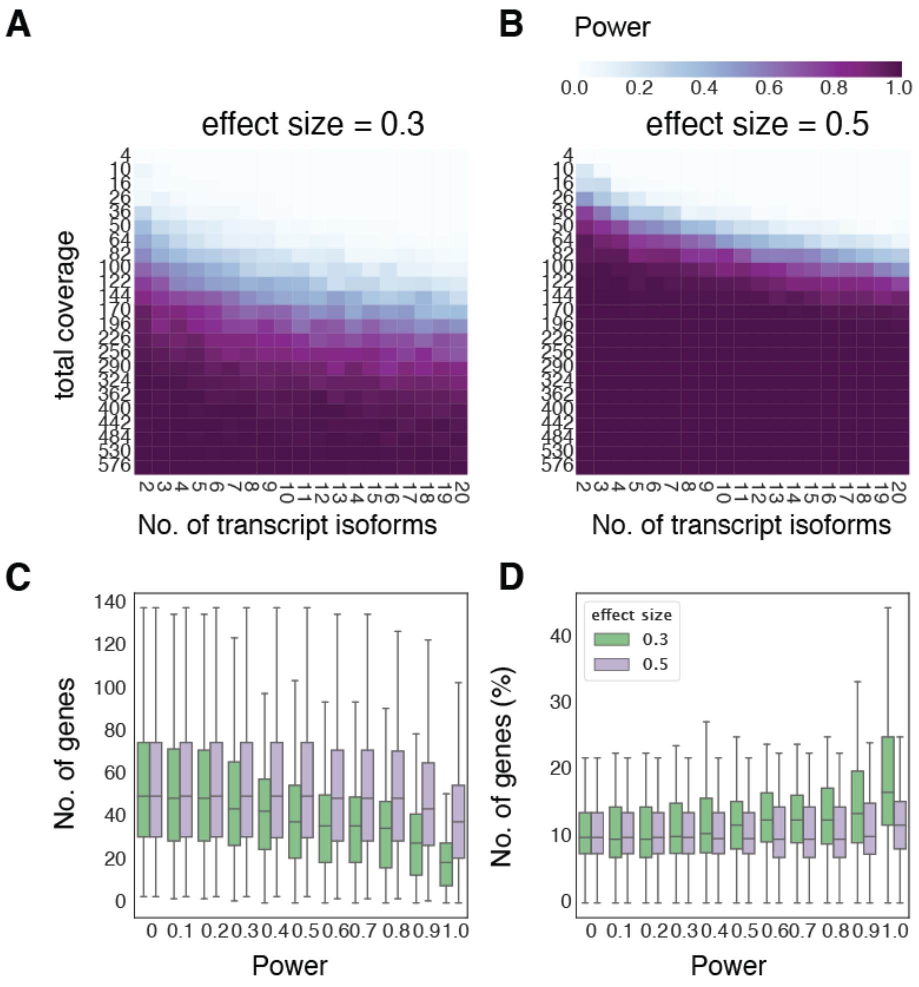


**Supplementary Figure 9:** Estimated power with respect to the number of transcripts (x-axis) and read counts (y-axis) for effect sizes **A)** 0.3 and **B)** 0.5, derived from simulated data. **C)** Total number of genes with more than one transcript, at different levels of power, including 67 samples processed at the Broad. **D)** The percentage of genes that have significantly different distributions (FDR < 0.05) of transcript expression in the two haplotypes, as a function of power. Outliers are hidden for ease of viewing.

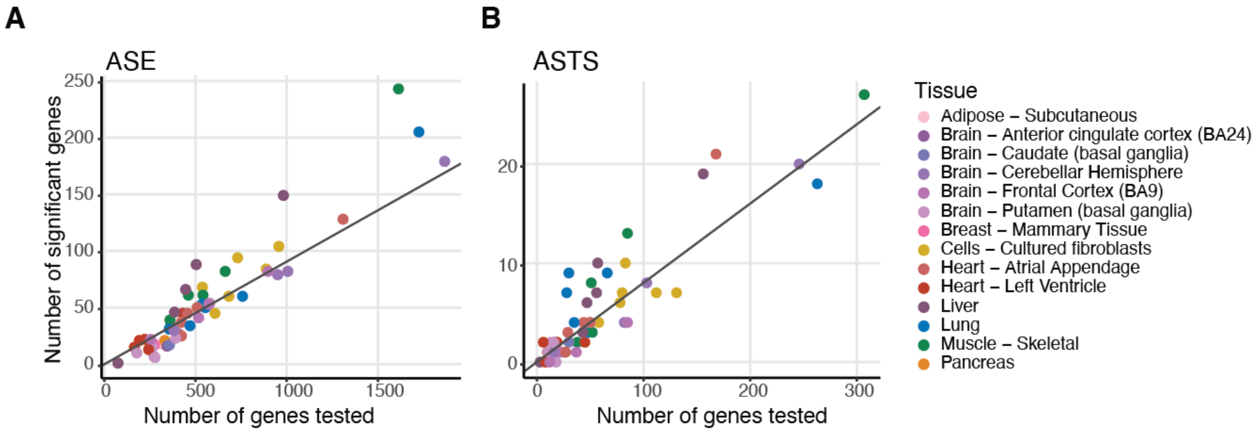


**Supplementary Figure 10**: Number of genes tested for **A)** allele-specific expression (ASE) and **B)** allele-specific transcript structure (ASTS) and number of significant genes. The diagonal indicate the median percentage of significant genes (9% and 9%, respectively).

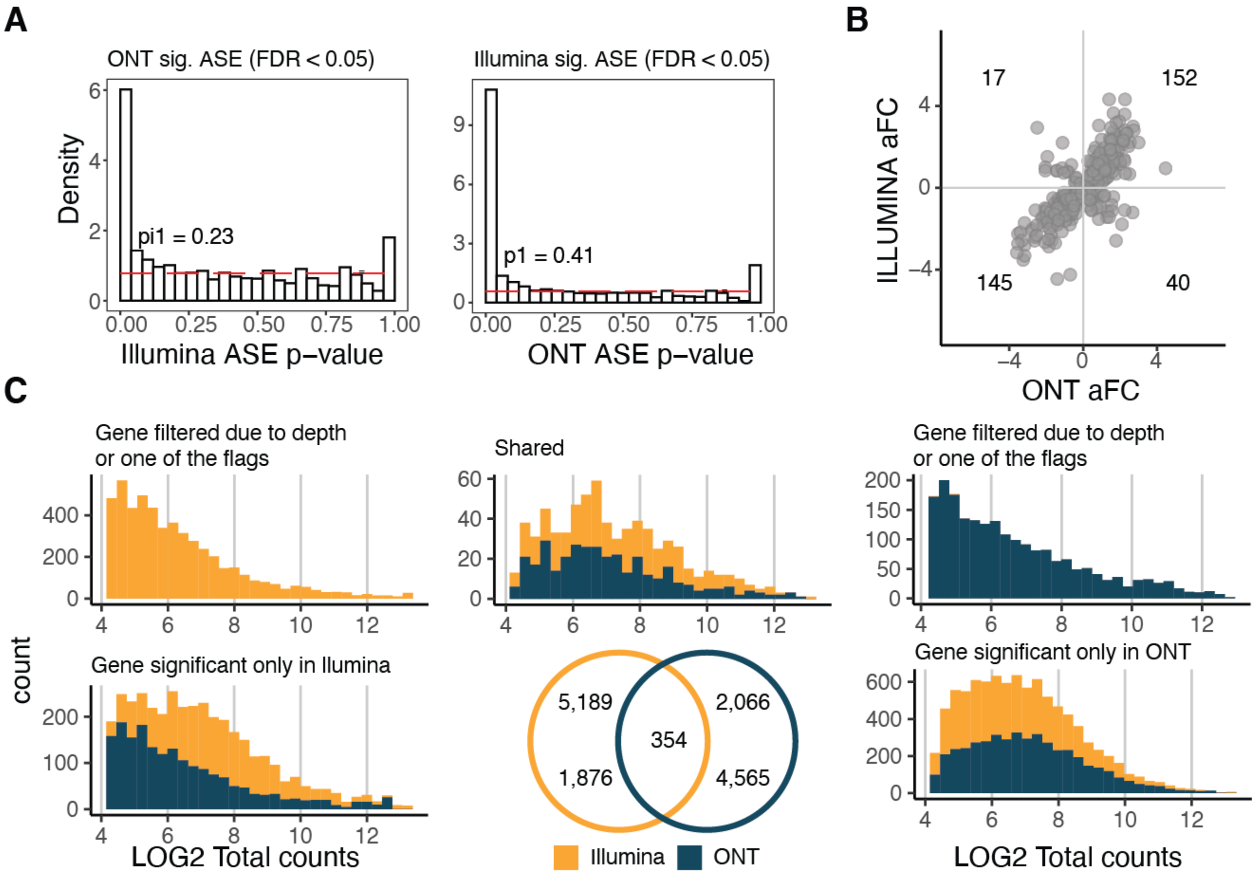


**Supplementary Figure 11**: **A)** Proportion of significant ASE genes discovered using Illumina or ONT and replicated in the other method. Pi1 calculations are carried out up to p-value = 0.5. **B)** Log allelic fold-change of Illumina and ONT of the shared ASE genes. **C)** Venn diagram of the significant ASE genes discovered by Illumina and ONT. The LOG2 of total counts for each method is shown for each group of the Venn diagram.

A

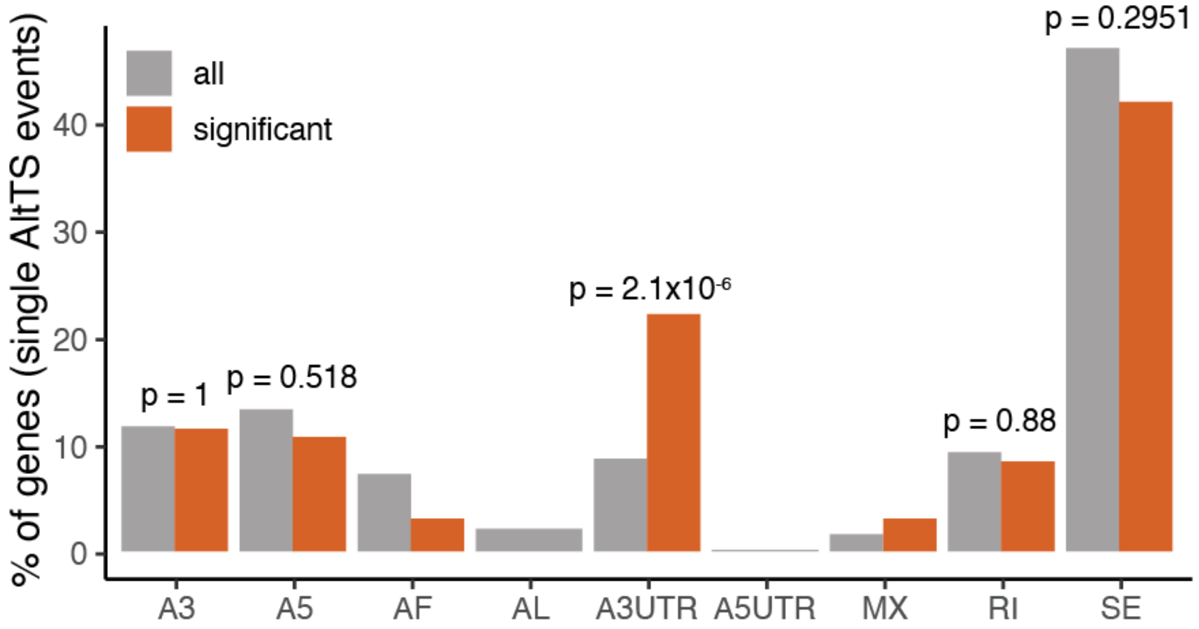


**Supplementary Figure 12**: **A)** Percentage of genes displaying a single alternative transcript structure event. A3: alternative 3' splice site; A5: alternative 5' splice site; AF: alternative first exon; AL: alternative last exon; A3UTR: alternative 3' end; A5UTR: alternative 5' end; MX: mutually exclusive exons; RI: retained intron; SE: skipped exon.

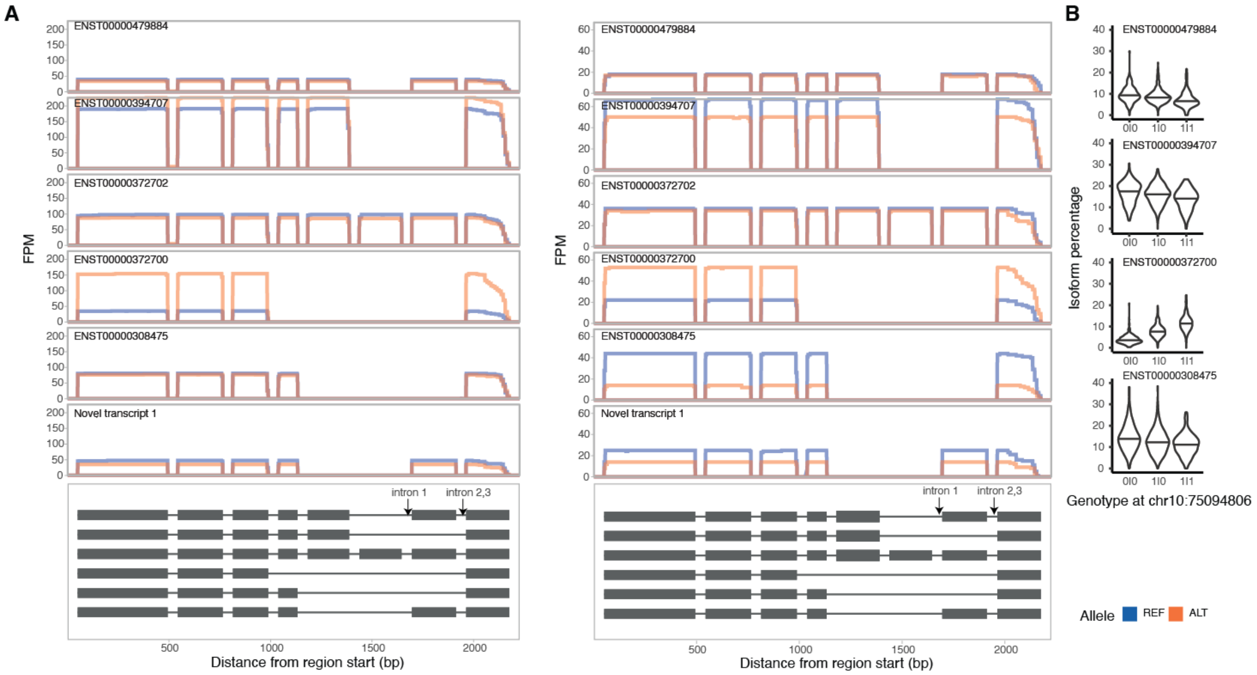


**Supplementary Figure 13**: **A)** Read pile-ups per transcript for the two donors displaying ASTS in *DUSP13* gene. In the lower panel the transcript structure is shown, without details of the coding sequence. **B)** Transcript percentage for four of the five annotated transcripts with high read coverage in the GTEx v8.

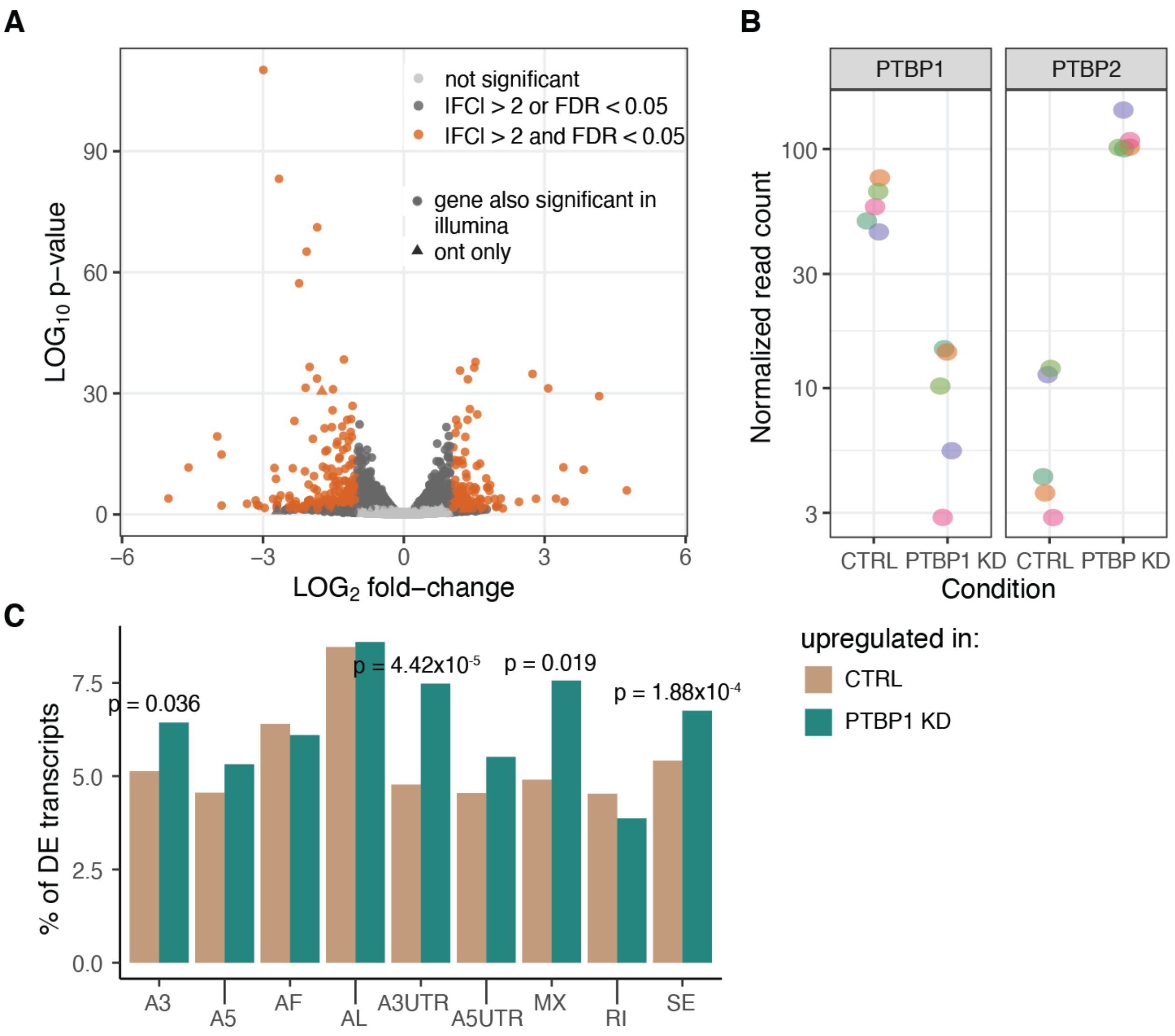


**Supplementary Figure 14**: **A)** Volcano plot from differential gene expression between control and samples with PTBP1 knockdown using ONT data. **B)** Gene expression profile in PTBP1 and PTBP2 genes (PTBP2 under normal circumstances has its expression restricted to the brain). **C)** Comparison between transcripts upregulated in the control or the PTBP1 knockdown samples across different alternative transcript structure events. P-values were calculated using a chi-square proportion test.

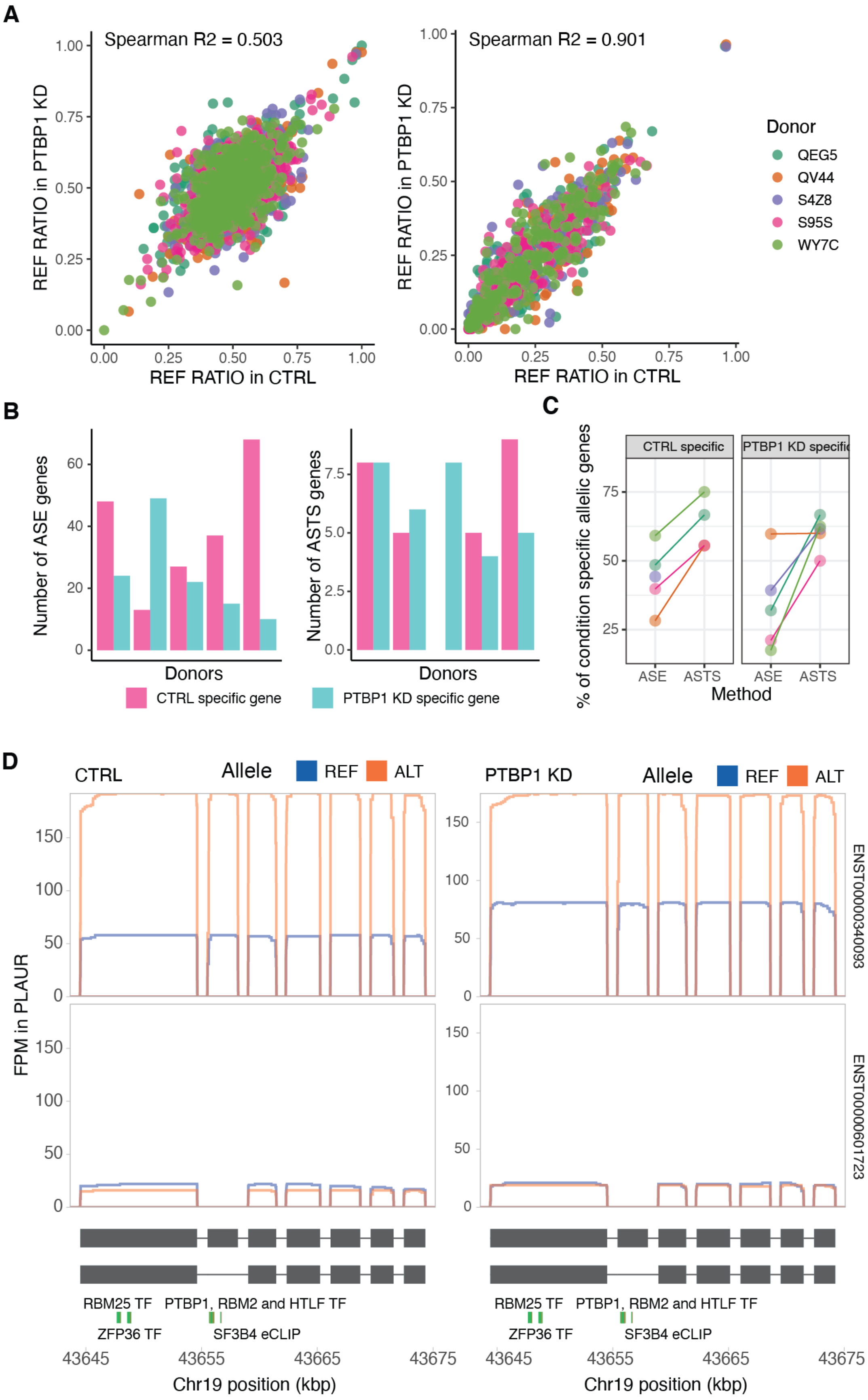

A
Spearman R2 = 0.503
REF RATIO in PTBP1 KD
REF RATIO in CTRL
Spearman R2 = 0.901
Donor
QEG5
QV44
S4Z8
S95S
WY7C
B
Number of ASE genes
Number of ASTS genes
Donors
CTRL specific gene
PTBP1 KD specific gene
C
% of condition specific allelic genes
CTRL specific
PTBP1 KD specific
ASE
ASTS
Method
D
CTRL
PTBP1 KD
Allele
REF
ALT
FPM in PLAUR
ENST00000340093
ENST00000601723
RBM25 TF
PTBP1, RBM2 and HTLF TF
ZFP36 TF
SF3B4 eCLIP
Chr19 position (kbp)

**Supplementary Figure 15**: **A)** Correlation between the control and the PTBP1 knockdown samples in the reference ratio of gene expression and transcript structure. **B)** Number of ASE and ASTS specific genes per condition across the five donors. **C)** Proportion of genes displaying allele-specific patterns specifically in either control or PTBP1 knockdown samples. **D)** PLAUR gene transcript read pile-ups which display significant ASTS only in the control sample. Also shown are the transcript structures and the eCLIP and ChIP-seq for RNA binding proteins as assessed by ENCODE. The skipped exon contains a RNA protein binding domain enriched in PTBP1, RMB2 and HTLF immunoprecipitation, within which the heterozygous variant is found.

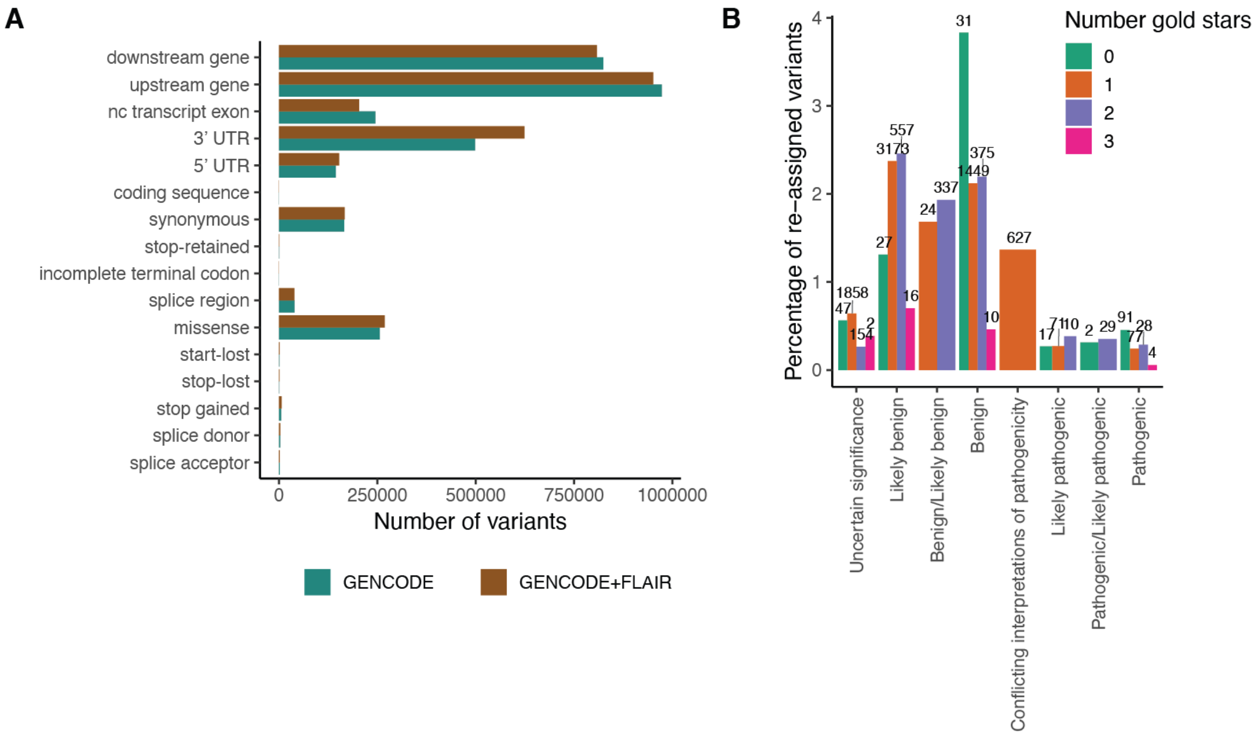


**Supplementary Figure 16**: **A)** Number of variants per variant effect predictor (VEP) category using GENCODE v26 protein-coding genes with or without novel FLAIR transcripts. **B)** Percentage of variants per clinical significance category that get reassigned when supplementing the gene annotation with the novel transcripts. The numbers above the bars correspond to the number of re-assigned variants.

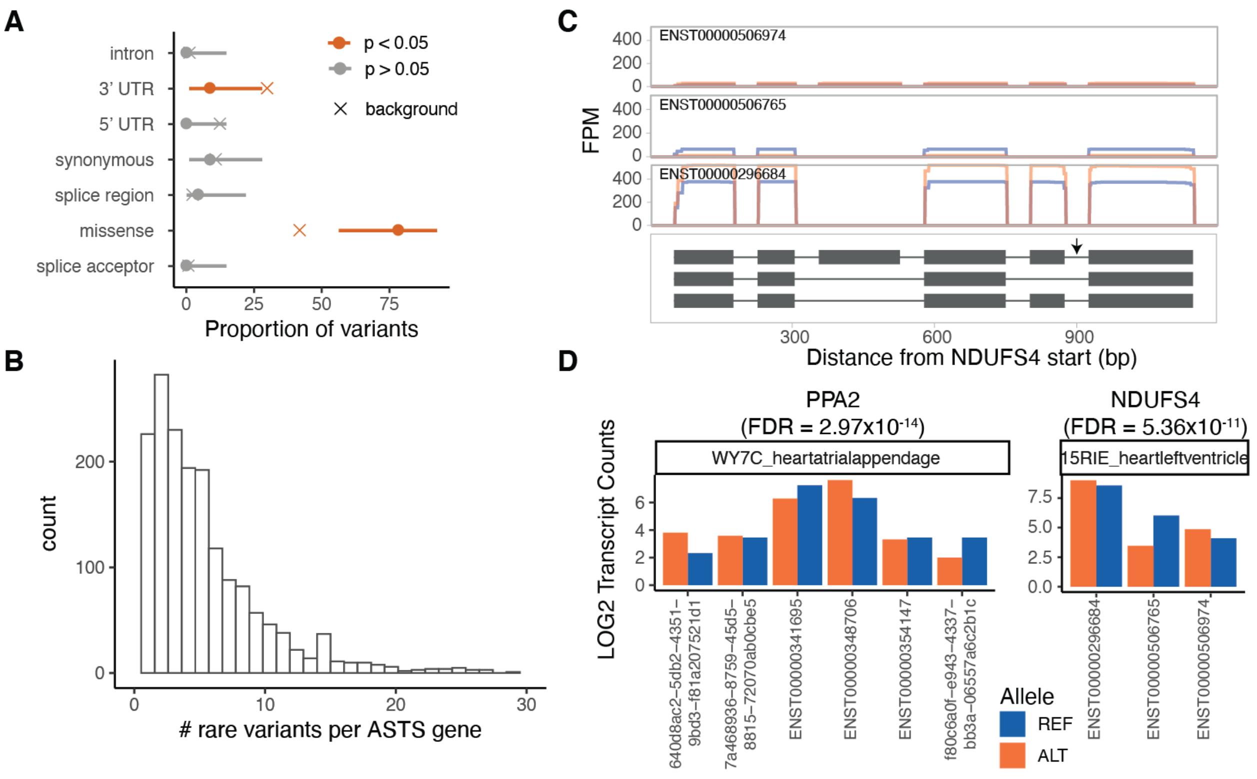


**Supplementary Figure 17**: **A)** Proportion of rare heterozygous variants per annotation in significant ASTS events. As a background all ASTS events were used, and p-values were calculated using binomial testing. **B)** Number of rare variants per ASTS gene (10kb window around gene). **C**) *NDUFS4* as an example of a gene with a rare heterozygous variant in a sample that is a GTEx splicing outlier and has significant ASTS, with read pileups, with grey arrows indicating the rare variants. **D)** Log normalised transcript counts per allele for *PPA2* and *NDUFS4*.

### References


1. Li, H. Minimap2: pairwise alignment for nucleotide sequences. *Bioinformatics* **34**, 3094–3100 (2018).
2. De Coster, W., D'Hert, S., Schultz, D. T., Cruts, M. & Van Broeckhoven, C. NanoPack: visualizing and processing long-read sequencing data. *Bioinformatics* **34**, 2666–2669 (2018).
3. Tang, A. D. *et al.* Full-length transcript characterization of SF3B1 mutation in chronic lymphocytic leukemia reveals downregulation of retained introns. *Nat. Commun.* **11**, 1438 (2020).

4. Workman, R. E. *et al.* Nanopore native RNA sequencing of a human poly(A) transcriptome. *Nat. Methods* **16**, 1297–1305 (2019).
5. Pertea, G. & Pertea, M. GFF Utilities: GffRead and GffCompare. *F1000Research* vol. 9 304 (2020).
6. Trincado, J. L. *et al.* SUPPA2: fast, accurate, and uncertainty-aware differential splicing analysis across multiple conditions. *Genome Biol.* **19**, 40 (2018).
7. Jiang, L. *et al.* A Quantitative Proteome Map of the Human Body. *Cell* **183**, 269–283.e19 (2020).
8. Keller, A., Nesvizhskii, A. I., Kolker, E. & Aebersold, R. Empirical statistical model to estimate the accuracy of peptide identifications made by MS/MS and database search. *Anal. Chem.* **74**, 5383–5392 (2002).
9. Deutsch, E. W. *et al.* Trans-Proteomic Pipeline, a standardized data processing pipeline for large-scale reproducible proteomics informatics. *Proteomics Clin. Appl.* **9**, 745–754 (2015).
10. Love, M. I., Huber, W. & Anders, S. Moderated estimation of fold change and dispersion for RNA-seq data with DESeq2. *Genome Biol.* **15**, 550 (2014).
11. Nowicka, M. & Robinson, M. D. DRIMSeq: a Dirichlet-multinomial framework for multivariate count outcomes in genomics. *F1000Research* vol. 5 1356 (2016).
12. Edge, P., Bafna, V. & Bansal, V. HapCUT2: robust and accurate haplotype assembly for diverse sequencing technologies. *Genome Res.* **27**, 801–812 (2017).
13. Castel, S. E., Levy-Moonshine, A., Mohammadi, P., Banks, E. & Lappalainen, T. Tools and best practices for data processing in allelic expression analysis. *Genome Biol.* **16**, 195 (2015).
14. Mohammadi, P., Castel, S. E., Brown, A. A. & Lappalainen, T. Quantifying the regulatory effect size of cis-acting genetic variation using allelic fold change. *Genome Res.* **27**, 1872–1884 (2017).
15. Cohen, J. *Statistical Power Analysis for the Behavioral Sciences*. (Academic Press, 2013).

16. GTEx Consortium. The GTEx Consortium atlas of genetic regulatory effects across human tissues. *Science* **369**, 1318–1330 (2020).

17. Van Nostrand, E. L. *et al.* A large-scale binding and functional map of human RNA-binding proteins. *Nature* **583**, 711–719 (2020).

18. Quinlan, A. R. & Hall, I. M. BEDTools: a flexible suite of utilities for comparing genomic features. *Bioinformatics* **26**, 841–842 (2010).

19. Gremme, G., Steinbiss, S. & Kurtz, S. GenomeTools: a comprehensive software library for efficient processing of structured genome annotations. *IEEE/ACM Trans. Comput. Biol. Bioinform.* **10**, 645–656 (2013).

20. Rentzsch, P., Witten, D., Cooper, G. M., Shendure, J. & Kircher, M. CADD: predicting the deleteriousness of variants throughout the human genome. *Nucleic Acids Res.* **47**, D886–D894 (2019).
